# Different historical generation intervals in human populations inferred from Neanderthal fragment lengths and patterns of mutation accumulation

**DOI:** 10.1101/2021.02.25.432907

**Authors:** Moisès Coll Macià, Laurits Skov, Benjamin Marco Peter, Mikkel Heide Schierup

## Abstract

After the main out-of-Africa event, humans interbred with Neanderthals leaving 1-2% of Neanderthal DNA scattered in small fragments in all non-African genomes today^1,2^. Here we investigate the size distribution of these fragments in non-African genomes^3^. We find consistent differences in fragment length distributions across Eurasia with 11% longer fragments in East Asians than in West Eurasians. By comparing extant populations and ancient samples, we show that these differences are due to a different rate of decay in length by recombination since the Neanderthal admixture. In line with this, we observe a strong correlation between the average fragment length and the accumulation of derived mutations, similar to what is expected by changing the ages at reproduction as estimated from trio studies^4^. Altogether, our results suggest consistent differences in the generation interval across Eurasia, by up to 20% (e.g. 25 versus 30 years), over the past 40,000 years. We use sex-specific accumulations of derived alleles to infer how these changes in generation intervals between geographical regions could have been mainly driven by shifts in either male or female age of reproduction, or both. We also find that previously reported variation in the mutational spectrum^5^ may be largely explained by changes to the generation interval and not by changes to the underlying mutational mechanism. We conclude that Neanderthal fragment lengths provide unique insight into differences of a key demographic parameter among human populations over the recent history.

## Introduction

If Neanderthal sequences in all non-Africans stem from a single introgression event, then differences in Neandertal fragment length distribution across the world would be indicative of differences in the speed of the recombination clock. Assuming a constant number of recombinations per generation, this would then imply differences in the number of generations since the admixture event and consequently differences in generation times among populations. While recent studies point towards a single gene flow event^2^, an additional admixture event private to Asians has also been proposed^6,7^ because Asian genomes carry larger amounts of Neanderthal sequence compared to European genomes. However, Asian genomes will also have more archaic fragments if a single gene flow common to Eurasians was followed by dilution of Neanderthal content in Europeans due to subsequent admixture with a population without Neanderthal admixture^2,8^.

An independent source of information for estimating differences in generation time is the rate and spectrum of derived alleles accumulating in genomes over a given amount of time^9,10^. Pedigree studies have shown that the yearly mutation rate slightly decreases when the generation time increases because the mutational burst in the germline before puberty represents a high proportion of new mutations in young parents^4^. Moreover, the relative proportion of different mutational types depends on both the paternal and maternal age at reproduction. This has been exploited to estimate differences in generation intervals for males and females between Neanderthals and humans^9^.

Here we investigate archaic fragment length distributions among extant non-Africans genomes from the Simon Genome Diversity Project (SGDP)^3^ and high coverage ancient genomes. We report strong evidence for a single Neanderthal admixture event shared by all Eurasian and American individuals, enabling us to make use of archaic fragment length distributions as a measure of generation intervals since admixture. Differences in estimated generation intervals are mirrored by concordant patterns of mutation accumulation, and suggest significant differences by up to 20% in the generation time interval experienced by different Eurasian regions since their splits.

## Results

### Neanderthal fragment length distributions differ across Eurasia

The average archaic fragment lengths in non-African individuals from the SGDP, inferred using the approach of Skov et al^11^, differs across Eurasia and America (Fig. 1a, S3, Data1_archaicfragments.txt). It presents a clear west-east gradient with the lowest mean fragment length in an individual from the Middle East (S_Jordanian-1, mean = 65.69 kb, SE = 2.49 kb, sd = 72.09 kb, S1) and the highest in an individual from China (S_Tujia-1, mean = 88.70 kb, SE = 3.29 kb, sd = 110.62 kb, S1). The pattern is qualitatively very similar when a) median fragment length instead of mean lengths are used, b) restricting to fragments most closely related to the Vindija Neanderthal genome, the sequenced Neanderthal that is most closely related to the introgressing Neanderthal population^12^ or c) only using high-confidence fragments inferred by the model (Extended Figure 1a-c, S4). When individuals are grouped into five main geographical regions, the average archaic fragment length distributions are significantly different (P value < 1e-5, permutation test, S2) by up to 1.12 fold (Fig. 1b zoom in, Table S1). These five regions also show significant differences in the number of archaic fragments and in the amount of archaic sequence inferred per individual (P value < 1e-5 for both, permutation test, S2, Fig. 1c and d, Table S1), mirroring the mean archaic fragment length distribution patterns. In agreement with previous reports^2,7^, we found, for example, that East Asians have 1.32 fold more archaic sequence inferred per individual compared to West Eurasians (P value < 1e-5, permutation test, S2, Fig. 1d, Table S1).

**Fig 1.**
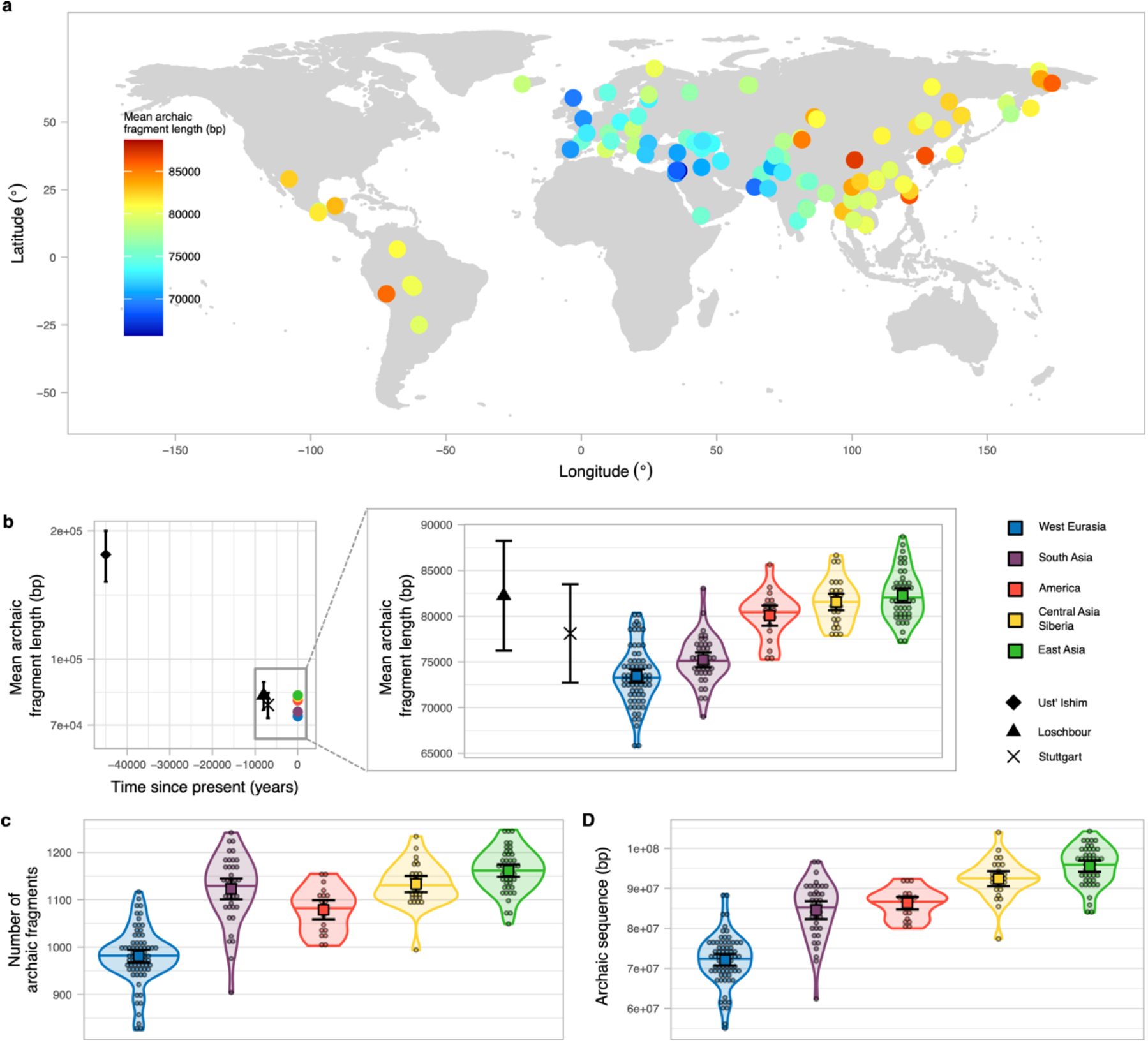
Archaic fragment statistics distributions around the world and in ancient samples. **a)** World map showing the samples from SGDP used in this study coloured according to the mean archaic fragment length. **b)** Mean archaic fragment length of extant geographical regions and ancient samples. Ust*’*-Ishim, Loschbour and Stuttgart mean archaic fragment length are shown as black points with specific shapes with their corresponding 95%CI as error bars. The average of the mean archaic fragment length among all individuals in each of the 5 main regions are shown as points (colour-coded). The zoom-in shows the mean archaic fragment length distribution per region (colour coded) as a violin plot. Individual values are shown as dots. The median is shown as a horizontal line in each violin plot. The mean and its 95%CI of each distribution is shown as a coloured square with their corresponding error bars. Loschbour and Stuttgart mean length are also shown for comparison. **c) and d)** the number of archaic fragments and the archaic sequence distributions respectively per region (colour coded) as violin plot. Individual values are shown as dots. The median is shown as a horizontal line in each violin plot. The mean and its 95%CI of each distribution is shown as a coloured square with their corresponding error bars. (width = 18cm)

We next investigated whether the larger amount of archaic sequence in East Asians is explained by having distinct archaic fragments due to a second Neanderthal admixture. We did this by joining the fragments of the 45 East Asian individuals and comparing them to the joined fragments of a subsample of 45 West Eurasian individuals (Extended Figure 2a and b, S6, Extended Figure 3). A total of 916,369 kb of the genome is covered by archaic sequence in East Asia and 866,945 kb in West Eurasia, with 485,255 kb (53%) of the archaic sequence overlapping (Fig. 2a, Table S3). Thus, as a group, East Asia has 5% more genomic positions with archaic introgression evidence. If we further remove fragments with the closest affinity to the sequenced Denisovan, which East Asians are known to possess more of^13^, the total sequence covered by archaic fragments is almost identical (East Asia 853,065 kb, West Eurasia 850,028 kb, Table S5). When we restrict to fragments with affinity to the Vindija or Altai Neanderthal, East Asia has a ∼7% higher proportion of the genome covered (East Asia 646,710 kb, West Eurasians 604,518 kb, Table S5). We ascribe this latter difference to the fact that shorter fragments in Western Eurasians both make them slightly harder to infer by the Skov et al^11^ approach and less likely to carry SNPs that directly classify them as closest to the Vindija Neanderthal.

**Fig 2.**
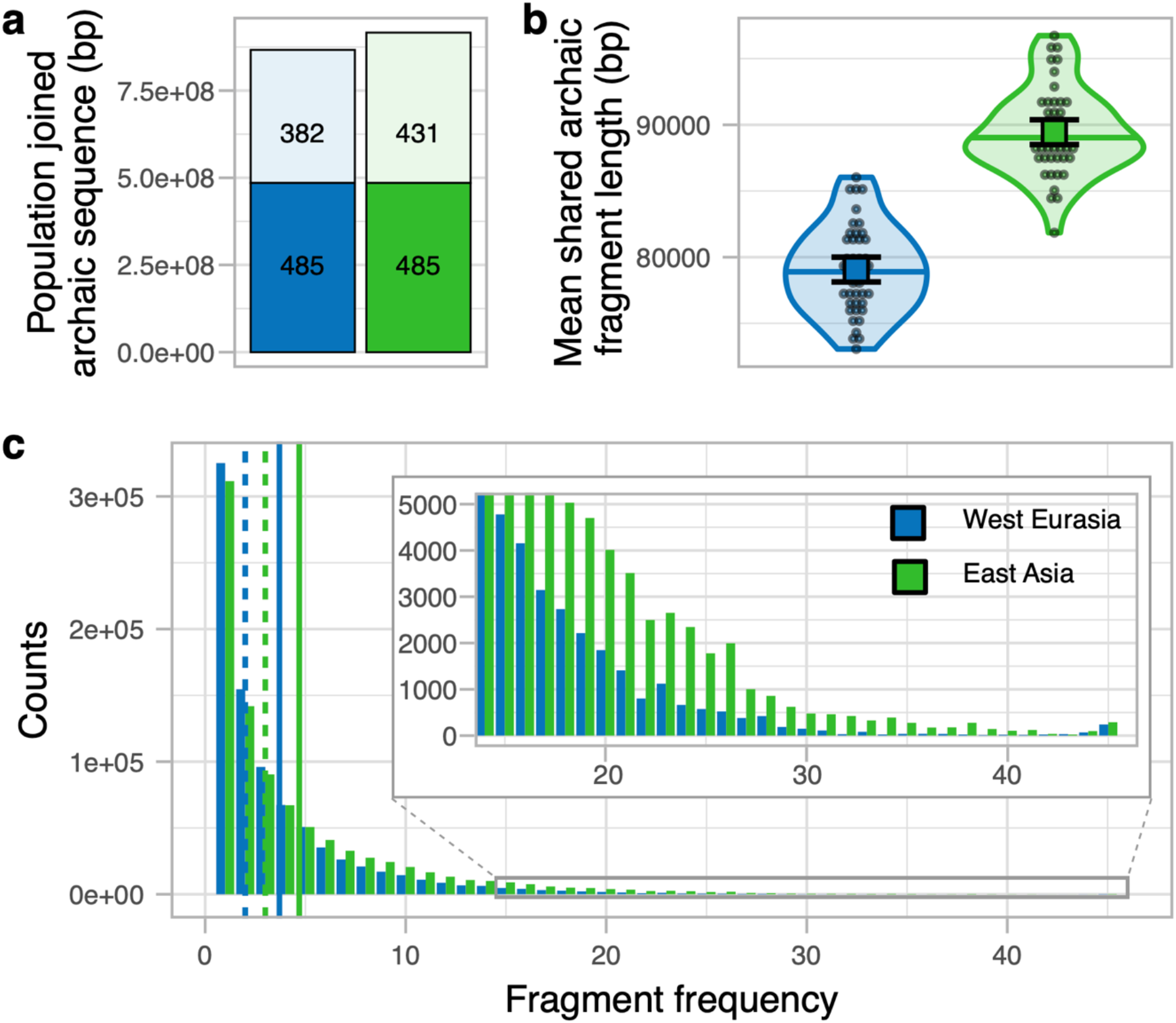
West Eurasia and East Asia archaic fragments comparison. **a)** Joined archaic sequence in both geographic regions (colour coded). The portion of the bar painted in plain colour shows the shared amount between the regions. The rest of the column shows the sequence private of each region. The numbers in each section denote the corresponding archaic sequence in Mb. **b)** The mean archaic fragment length distributions of individual shared fragments among regions per region (colour coded) as violin plot. Individual values are shown as dots. The median is shown as a horizontal line in each violin plot. The mean and its 95%CI of each distribution is shown as a coloured square with their corresponding error bars. **c)** The number of 1 kb genomic windows (y-axis) in which an archaic fragment has been found in a certain amount of individuals (x-axis) for each region. The insert shows the high-frequency bins. Vertical lines show the mean (plain lines) and median (dashed lines) for each region. (width = 9cm)

To compare shared fragments in terms of length, we only consider fragments in an East Asian that overlap with regions in the genome of West Eurasians that contain archaic sequence and vice versa (Extended Figure 1d, Extended Figure 2c, S6). We observe that shared fragments in East Asian individuals are on the average 1.13 fold longer than in West Eurasians (P value < 1e-5, permutation test, S2, Fig. 2b, Table S4) as also observed when all fragments were used.

Based on these observations, we conclude that the vast majority and possibly all of Neanderthal ancestry in East Asians and West Eurasians stems from the same Neanderthal admixture event. The 32% higher total amount of archaic sequence in an East Asian compared to a West Eurasian individual on average is primarily due to archaic fragments occurring at higher frequency in East Asians (Fig. 2c, Extended Figure 2d, S6, Extended Figure 3). It is unlikely that natural selection has acted much more strongly against archaic fragment frequency in West Eurasia since the purging of Neanderthal introgression is expected to have acted prior to the split of European and Asian populations^14,15^. We consider our observations more compatible with Europeans mixing with a Basal Eurasian population with little or no archaic content diluting the Neanderthal ancestry as has previously suggested from admixture modelling using ancient samples^8^. Such a dilution process should have a negligible effect on the Neanderthal fragment lengths observed today (Fig. 2b) but would shift the frequency distribution of Neanderthal fragments as we observe (Fig. 2c). This leaves us with a difference in the speed of the recombination clock and hence the number of generations since the common admixture with Neanderthals as the major cause of differences in archaic fragment length distributions.

Ancient genomes allow us to look at archaic fragment lengths back in time. We called archaic fragments in three high-coverage ancient samples included in the SGDP data; Ust*’*-Ishim (dated 45,000 BP, equally related to all Eurasians)^16^, Stuttgart (dated 7,000 BP farmer, West Eurasian ancestor)^17^ and Loschbour (dated 8,000 BP hunter-gatherer, West Eurasian ancestor)^17^ (S3, Fig. S1, Table S2). As expected, the archaic fragments are much longer for Ust*’*-Ishim compared to any of the ancient and extant individuals (Fig. 1b, Fig. S1, Table S2, see also ^16,18^). Loschbour*’*s and Stuttgart*’*s archaic fragments are on average longer than their West Eurasian descendants. However, their mean fragment length are very similar to the other extant populations, particularly East Asian populations (Fig. 1b zoom in) suggesting East Asians archaic fragments have experienced the same amount of recombination as West Eurasian ancestors, represented as Loschbour, 8,000 years ago. This corresponds to around 275 fewer generations in East Asia than in West Eurasian populations (assuming an average generation time of 29 years) over the approximately 40,000 years since the split of European and Asian populations^2,19–21^. Another way of stating this is that the 8,000 fewer years of recombination over 40,000 years corresponds to a difference in generation time of about 20% across Eurasia.

### Mutations accumulated differently across Eurasia

The number of *de novo* mutations (DNM) transmitted to a child depends on the sex and the age of the parents^4^. Thus, a change in generation time during recent human evolutionary history, as suggested above, should leave a detectable pattern in the total number of mutations accumulated. To test this, we estimated the number of derived alleles accumulated in each individual*’*s autosomes since the split of African and non-Afrcan populations (S7, Data2_mutationspectrum.txt). This was done by first removing all derived alleles observed in the Sub Saharan Africa outgroup, excluding those individuals with detectable West Eurasian ancestry^3^. Furthermore, we masked all genomic regions with evidence of archaic introgression in any individual since archaic variants would not be found in Sub-Saharan genomes and they would affect our results because they accumulated under a different mutational process^9^. Masking those regions also ensures that this analysis is independent of the archaic fragment length analysis above. After these procedures, we were left with ∼20% of the callable genome (S7).

Fig. 3a shows that the rate of accumulation of derived alleles is significantly different among groups (P value = 2.8e-4, permutation test, S2, Table S6). West Eurasia has accumulated 1.09% more derived alleles than East Asia (P value = 1.18e-3, permutation test, S2) since the Out-of-Africa event. However, this difference in the accumulation of derived alleles could only have happened when West Eurasia and East Asia were separated, which is only a part of the time since the Out-of-Africa (Fig. S3). If we assume >60,000 years for the out-of-Africa and a West Eurasia/East Asia split of <40,000 years^2,19–21^ the difference in the rate of derived allele accumulation is at least 60,000/40,000*1.09%=1.64% while West-Eurasia and East Asia were apart (SI8). Using the pedigree-based estimate of the relationships between mean parental age and mutation rate per generation^4^ (SI8), we estimate that this difference corresponds to a 2.68 or 3.39 years shorter generation interval in West Eurasia if East Asian mean generation time was 28 or 32 years respectively (SI8). These are lower bounds of the inferred differences in generation intervals since the difference between out-of-Africa and population split times is minimized.

**Fig 3.**
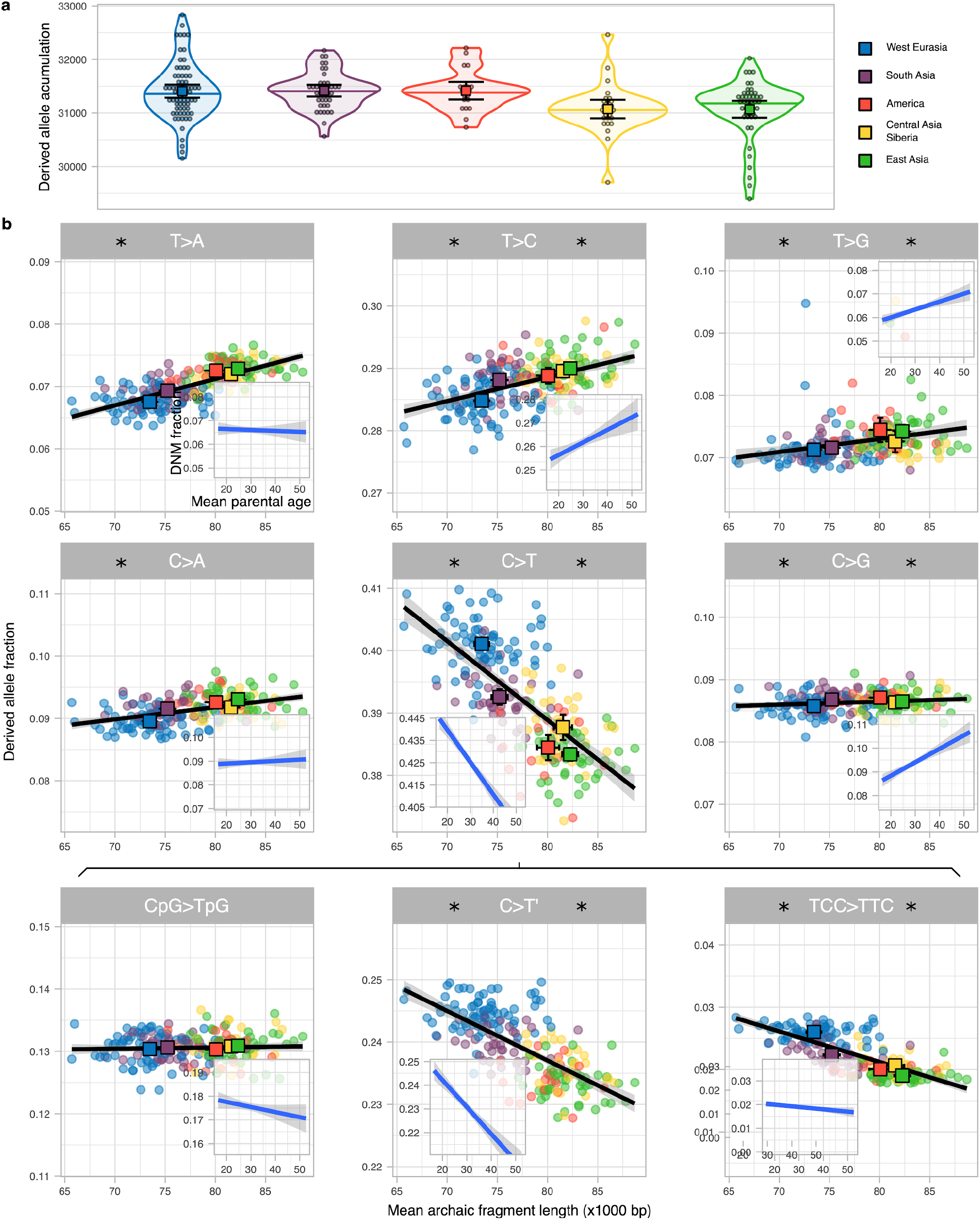
Derived allele accumulation distributions and their mutation spectrum. **a)** Distribution of the derived allele accumulation (y-axis) per region (colour coded) as violin plot. Individual values are shown as dots. The median is shown as a horizontal line in each violin plot. The mean and its 95%CI of each distribution is shown as a coloured square with their corresponding error bars. **b)** Correlation between the derived allele proportion (y-axis) with the mean archaic fragment length (x-axis) for 9 mutation types. Each dot represents an individual coloured according to the region they belong to. For each region, The mean and its 95%CI of both axes is shown as a coloured square with their corresponding error bars. Linear regressions (black lines) are shown with their corresponding SE (shaded area). For each mutation, the linear regression and corresponding SE between the fraction of DNM and mean parental age per proband of the deCODE data (**S9**) is shown as an insert. Note that the total span of the y-axis is the same for all panels and inserts but centred at the mean value specifically in each panel and insert. Asterisk on the left and right side of each mutation type indicates that the slope of the linear regression is significantly different from 0 for the SGDP and the deCODE data respectively. (width = 18cm)

The age of parents at conception, and hence generation time, also impacts the frequency of which types of single nucleotide mutations occur^4^. Thus, a shift in generation time is predicted to change the spectrum of new mutations^9,10^ and partially explain differences in mutation spectrum described among human populations^5,22,23^. We calculated the relative frequencies of the six different types of single nucleotide mutations depending on their ancestral and derived allele (S7, Fig. S2, Table S7) and related that to the average Neanderthal fragment length for each individual (Fig. 3b). We observe significant associations with average archaic fragment lengths for all six types (Table S8). We further subdivide C>T mutations in three types: CpG>TpG which present a distinct mutational process^24^ (Fig. S2, Table S7), TCC>TTC, which is in great excess in European genomes and has been studied as a population-specific mutational signature^5,22^ (Fig. S2, Table S7) and the rest, denoted as C>T*’* (Fig. 3b, Table S8). We find that the frequency of CpG>TpG transitions depends the least on fragment length.

To investigate whether these correlations could be due to differences in generation time between geographical regions, we reanalysed the proportion of DNM mutation types as a function of mean parental ages in the deCODE trio data set^4,25^ (Fig. 3b inserts, S9, Table S8). Comparing the correlations from the SGDP data with the deCODE data we see a strong correspondence for most mutational types: in all mutation types where correlations with either dataset are significant, the direction of the effects are concordant (Figure 3b, Fig. S6). The deCODE dataset has a slight bias towards probands having older fathers than mothers (mean = 2.77 years, sd = 4.25, Fig. S5), and this could affect the response of mutation type fraction depending on mean parental age. However, no major change in the correlation coefficients was observed when only probands with similar parental ages were analysed (S9, Fig. S6).

Since there is no a priori reason to expect a relationship between archaic fragment lengths and derived allele accumulation, we consider it likely that the same underlying factor has affected both. The general correspondence of these correlations with those expected from DNM studies supports our hypotheses that this causal element is a change in generation time. More specifically, the matching decreasing correlation with parental age of TCC>TTC mutation indicates that this mutation signature will increase when the mean parental age decreases. Thus a considerable reduction in mean generation time in West Eurasians, as suggested in this study, offers an alternative explanation to the excess of TCC>TTC mutations in that region compared to the rest of the world^5,26^.

An increase in the mean generation interval can be due to an increase in paternal or maternal age, or both. Anthropological studies suggest that males have generally been older than females at reproduction, but that the age gap is twice as large in hunter-gatherers compared with sedentary populations^27^. To gain insight into sex-specific changes to generation time intervals we first compared the accumulation of derived mutations between autosomes, which spend the same amount of evolutionary time in both sexes, and X chromosomes, which spend ⅔ of the time in females while ⅓ in males (S10). Thus, an increase of the male-to-female generation interval is expected to increase the X chromosome to autosomes (X-to-A) mutation accumulation ratio^28^, although other factors such as reproductive variance and changes in population size can also influence the ratio. Fig. 4a shows the X-to-A ratio of derived alleles accumulated per base pair as a function of the mean archaic fragment length, as mean generation time proxy, for the females in the SGDP data (S10). We observe that the X-to-A ratio is significantly different among regions (P value = 3.6e-4, permutation test, S2). East Asians have a higher X-to-A ratio compared to American and Central Asia and Siberia, with similar Neanderthal fragment sizes, and higher than West Eurasians, with smaller Neanderthal fragment sizes. This result is compatible with East Asians having a higher mean generation time than West Eurasians primarily due to an increased paternal age at reproduction as compared to Americans and Central Asia and Siberia where the age at reproduction of both sexes are inferred to have increased similarly.

**Fig 4.**
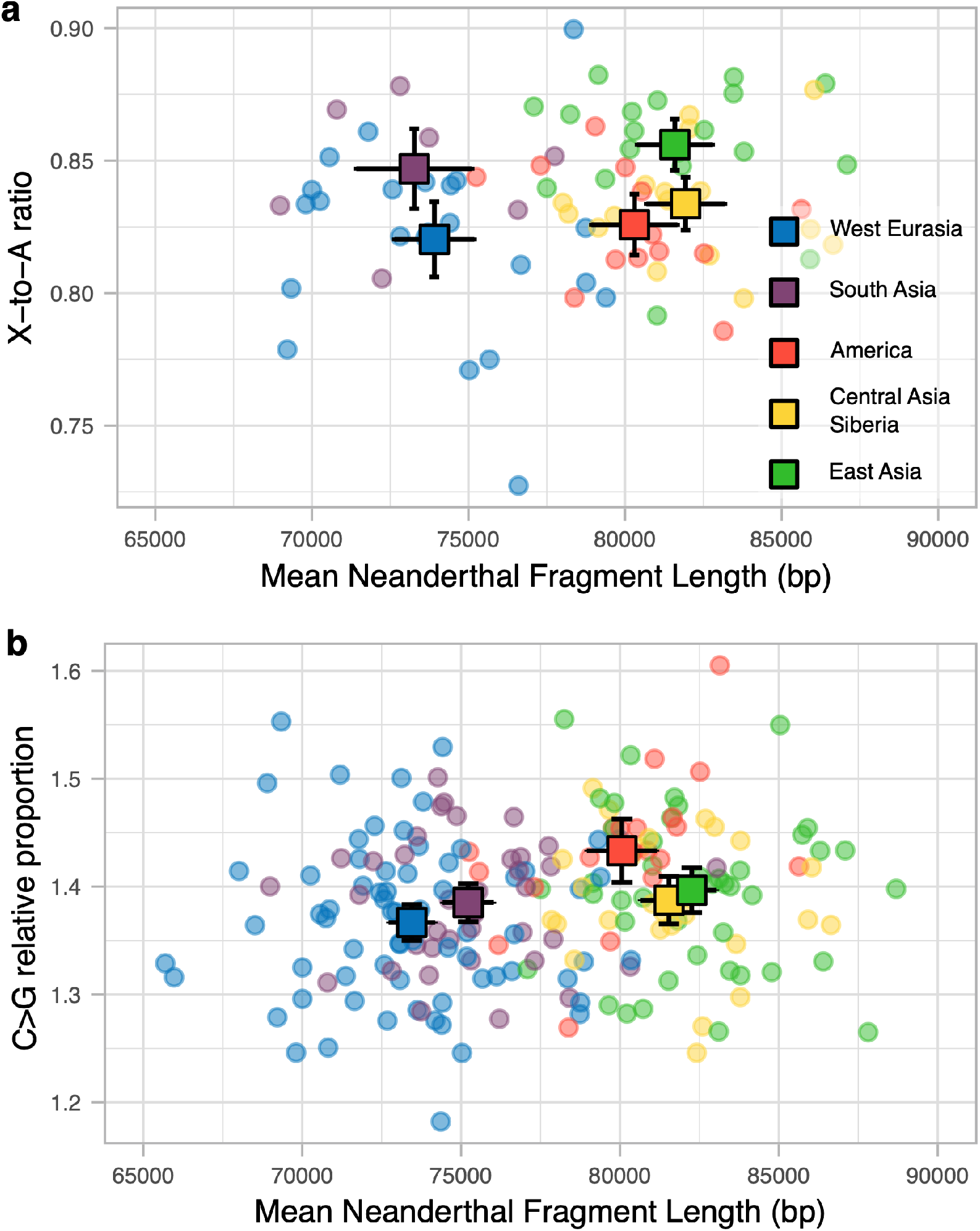
Sex-specific mutation patterns. **a)** Scatterplot of the X chromosome vs Autosome derived allele accumulation ratio (y-axis) and the mean Neanderthal fragment length (x-axis) for each region (colour coded). Each dot represents an individual in the corresponding population. The mean for each population for each axis is shown in squares and their 95%CI are denoted by the error bars. Only females were used to produce this plot. **b)** The same as in **a**, but the ratio between the proportions of C>G derived alleles in cDNM and the rest of the genome (**S10**). All samples were used to produce this plot. (width = 9cm)

Another sex-specific mutation signature are C>G mutations in genomic regions with clustered de novo mutations in old mothers^4,29^. This signature can be explored to compare maternal ages among groups^9^. We estimated the proportion of derived C>G alleles to other derived allele types in these genomic regions and contrasted it to the same ratio for the rest of the genome, for each individual (S10). When samples are grouped in the 5 main regions, the C>G ratio in DNM clusters differs significantly (P value = 2.77e-3, permutation test, S2), increasing with increasing Neanderthal fragment length (Fig. 4b). Notably, America has a higher ratio than Central Asia and Siberians for similar Neanderthal fragment lengths, suggesting a relatively larger impact of old mothers to the overall mean generation time throughout their history. This is in line with the X chromosome analysis in that longer generation times in America were more driven by older mothers as compared to older fathers in East Asia with an intermediate increase of both parental ages in Central Asia and Siberia.

Finally, the Y chromosome is also expected to accumulate more derived alleles in populations with younger fathers, similarly to the autosomes, about 0.4-0.5% per year difference in generation time between two populations. We observe a point estimate of 1.19% larger accumulation between West Eurasia and East Asia (Fig. S7, Table S10) but this is not significant with the limited data available for the Y chromosome (P value = 0.66, permutation test, S2).

## Discussion

We have shown that the length of Neanderthal fragments in modern human genomes can be used to obtain meaningful information about a fundamental demographic parameter, the mean generation interval. We estimate surprisingly large differences across eurasian and american groups suggesting stable differences over tens of thousands of years. Our approach depends on the assumption that archaic fragments trace back to a single Neanderthal admixture event shared by all non-African populations, for which we provide further evidence. Consistent with these results, the number of derived mutations accumulated in the geographic regions studied here follow the expectations of the difference in generation time estimated from the fragment lengths. The agreement between the recombination and the mutation clock signatures argues against confounding factors. For example, a potential bias would be expected if the African outgroup, here used to find archaic fragments in the other individuals, had experienced some ancient gene flow from West Eurasia that we have not been able to detect. Such a scenario would shorten and remove archaic fragments in West Eurasians, explaining the observed gradient. However, it would also decrease the number of derived alleles in West Eurasia compared to East Asia, which is the opposite to what we report.

Differences in generation intervals of the magnitude and duration that we estimate can account for observed variation in the mutation spectrum of human populations without an underlying change to the mutational repair system. An example of this is the increased frequency of the TCC>TTC mutation in West Eurasians. The differences in generation time, inferred here from archaic fragment lengths, explain more than half of the total variation among individuals (adjusted R^2^ = 55.53%).

Our results have direct implications for previous investigations of demographic human parameters, which have typically assumed that the generation interval was shared and constant for distinct human populations. Thus, future investigations should take variation in the generation time under consideration. We do not have an explanation for the underlying causes of large generation interval differences, but it is plausible that both low population densities and harsh environmental conditions increase generation time, whereas agriculture decreases mean generation times and reduces generation time differences between sexes. With an increasing number of sequenced ancient and modern genomes we anticipate that the approach we present here can be used to obtain a fine-grained picture of shifts in generation interval during the last 40,000 years that can be directly related to changes in population densities, climate and culture.

## Methods

A description of all analyses performed in this study is detailed in the Supplementary Information.

## Data availability

The archaic fragments and their basic statistics are provided in Data1_archaicfragments.txt; the counts of the 96 mutation types per individual per chromosome are provided in Data2_mutationspectrum.txt (S11).

## Code availability

The scripts coded to produce data and tables, perform statistical analysis and plot figures for this manuscript are accessible on Github (https://github.com/MoiColl/TheGenerationTimeProject).

## Acknowledgements

We thank Felix Riede for advice on anthropological interpretations and Matthew Hurles for suggesting to contrast derived allele accumulation between X and autosomes. We thank Juraj Bergman and Marjolaine Rouselle for long and fruitful discussions about the consequences of the Neanderthal dilution scenario in West Eurasian populations. We thank Priya Moorjani for reviewing and commenting on the study and giving insightful suggestions. The study was supported by grants NNF18OC0031004 from the Novo Nordisk Foundation and 6108-00385 from the Research Council of Independent Research to M.H.S.

## Contributions

M.C.M., L.S. and M.H.S designed the study. M.C.M. and L.S. created the methods to assess the data and, with M.H.S., analysed the results with input from B.M.P. M.C.M., L.S. and M.H.S. wrote the manuscript with comments from B.M.P.

## Competing interests

The authors declare no competing interests.

**Extended Figure 1.**
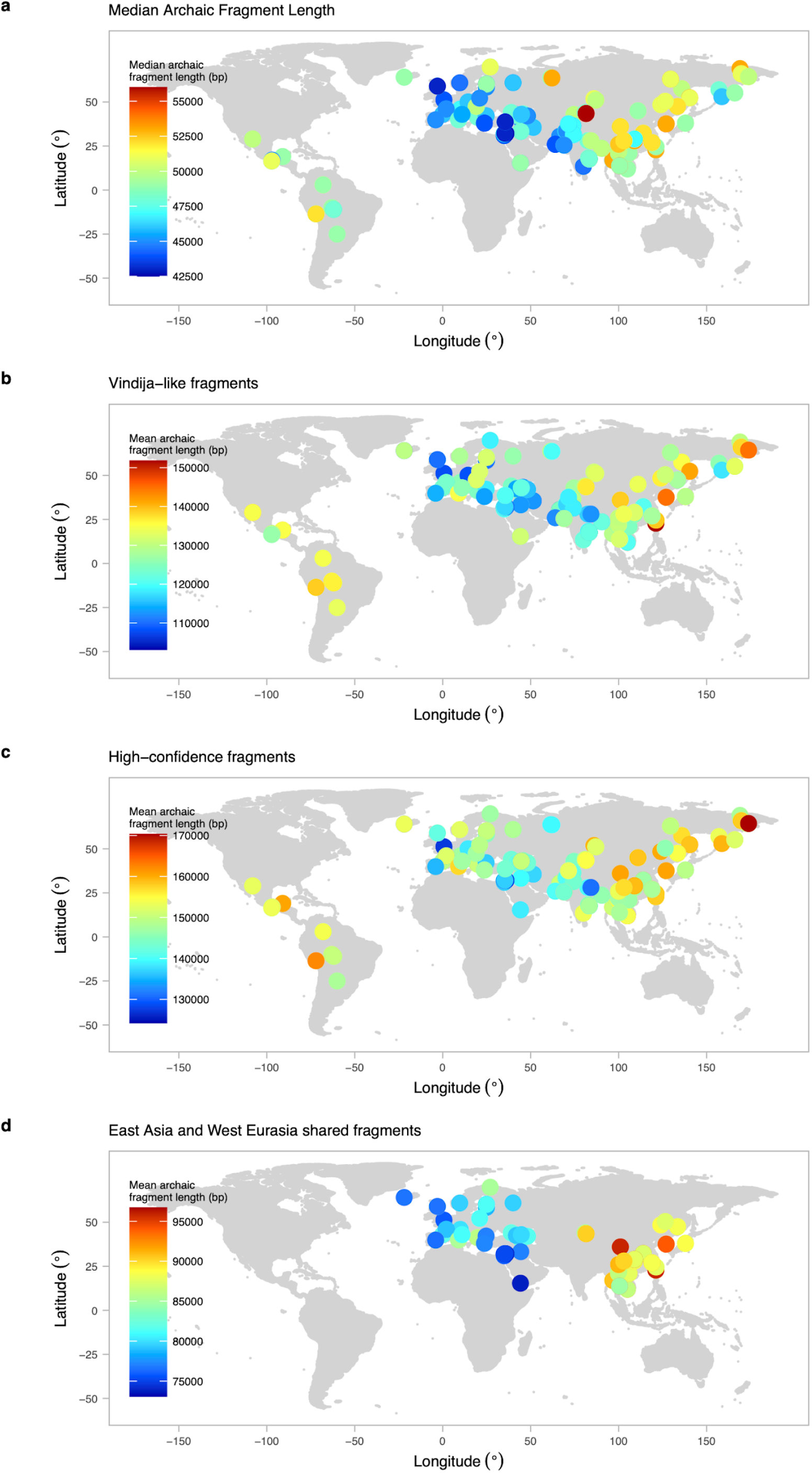
Archaic fragment length distribution around the world with specific filters. World map with samples from SGDP used in this study coloured according to the mean or median average archaic fragment length applying filters to the data. **a)** Median archaic fragment length is plotted instead of the mean. **b)** Only fragments with more SNPs shared with the Vindjia genome than the Denisova or the Altai genomes are used **c)** Only high confidence archaic fragments (posterior probability >= 90%) are used. **d)** Only shared individual fragments (Extended Figure 2, **S6**) between East Asians and West Eurasians.

**Extended Figure 2.**
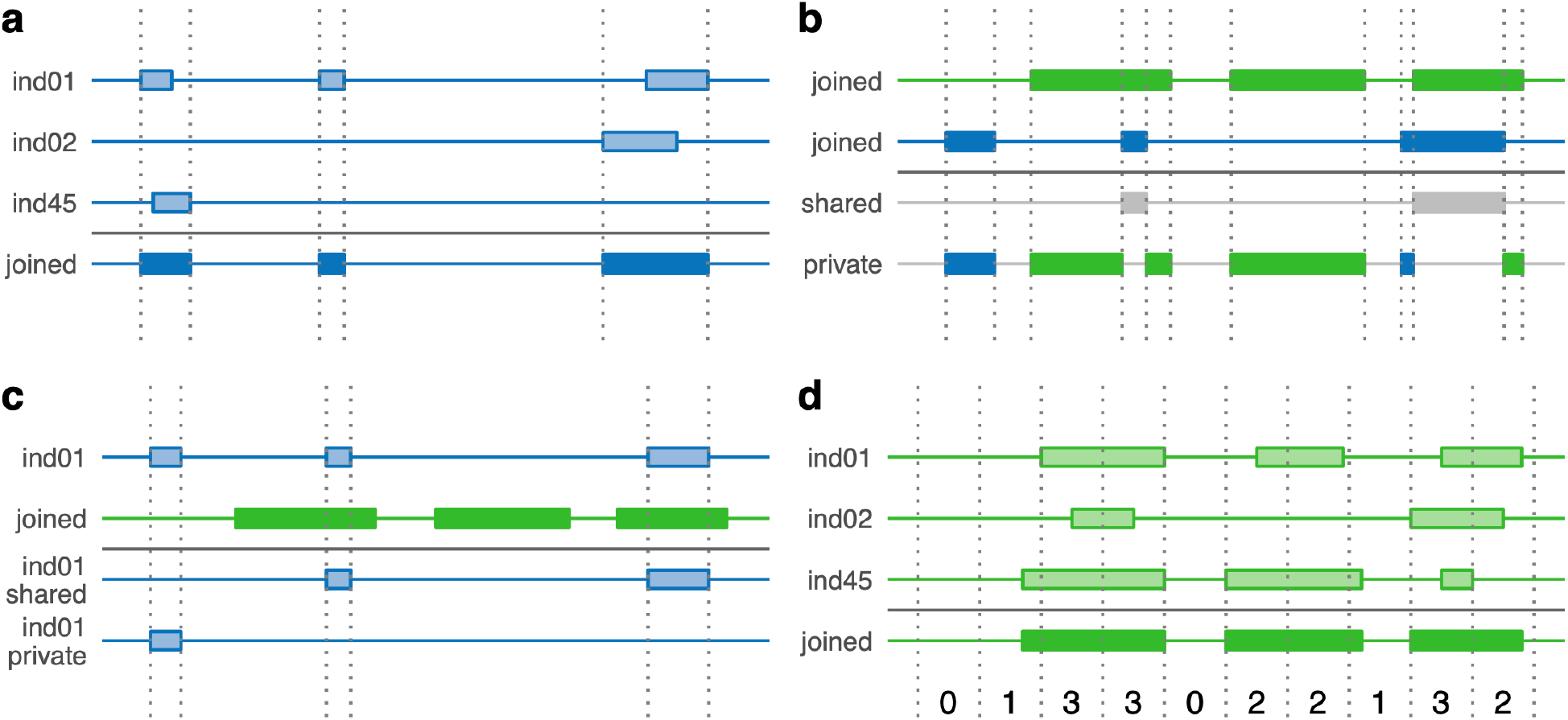
West Eurasia and East Asia fragment comparison methods. Diagram showing the different methods to compare archaic fragments between West Eurasians and East Asians (**S6**). Each horizontal line represents a genome. Wide bands on each genome represent archaic sequences. East Asia is represented in green colours and West Eurasia in blue. Grey colours are used when sequences are shared by both. Plain colours denote joined sequence and transparent colours show individual sequences. Vertical dashed lines are mainly used to point to genomic windows of interest. **a)** Joined region fragments. **b)** Shared and private joined region sequence. **c)** Shared and private individual fragments. **d)** Archaic frequency in 10 kb windows represented as the vertical grey lines intervals (note that in the main text, 1 kb windows are used instead).

**Extended Figure 3.**
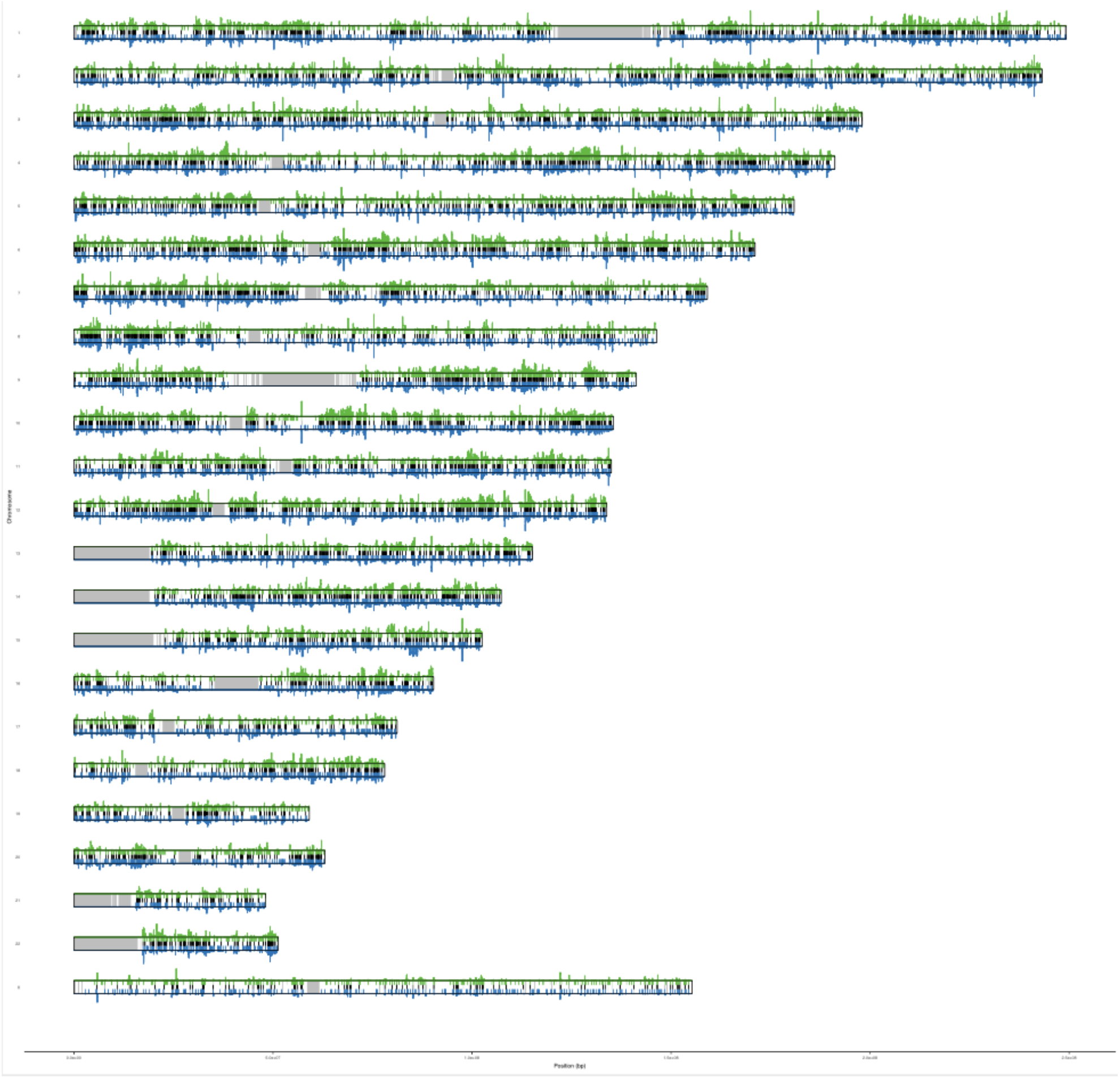
The archaic landscape across the West Eurasian and East Asian genomes. Each horizontal rectangle represents a chromosome (hg19). In each chromosome, it is shown the joined region fragments for West Eurasia (blue upper bands) and East Asia (green lower bands). The shared joined region fragments are shown as black bands in the middle of each chromosome. For each region, the number of individuals that have an archaic fragment in a particular 1kb window are represented as lines (maximum number of individuals is 45 for each region). Grey bands on the chromosomes show the non-callable portions of the genome (hg19).

## Supplementary Information

### 1 Confidence Interval calculation

The mean and its confidence intervals (CI) for any statistic are calculated using the mean and standard deviations of the 100,000 bootstrap sampling distribution of the observed statistic. The code to compute them is provided on the GitHub page.

### 2 Statistical significance assessment by permutation test

The statistical significance of a statistic to compare different groups is assessed by contrasting the observed statistic with a non-parametric null distribution. The null distribution is generated by permuting 100,000 the original data and calculating the statistic in each permutation. P values are then calculated as the fraction of permutations which yield a value as extreme or more extreme than what is observed in the data. If no such event is observed in all permutations, we considered the fraction to be < 1/100,000 = 1e-5. The significance level (α) in all tests is considered to be 0.05.

To test if there are differences between two groups for a statistic (for example, average archaic fragment length), we subtract the means of each group. In this case, since this test is a two-tailed hypothesis test, we multiply the obtained P value by two. When we test differences for multiple populations, we compute the F statistic.

The code to compute the statistical significance is provided on the GitHub page.

### 3 Identification of archaic fragments in non-African individuals and ancient samples

We called archaic fragments in individuals of the Simon Genome Diversity Project (SGDP) from West Eurasia, South Asia, America, Central Asia Siberia and East Asia regions as described in ^9,11^ -a step by step tutorial is also available at https://github.com/LauritsSkov/Introgression-detection.

In short, the method first removes a set of variants (SNPs) which are present in an outgroup with no presumed archaic admixture (Sub-Saharan African populations) from the samples in which we want to detect archaic fragments (non-Africans). Then, taking into account window-specific mutation rate and callability, the method classifies non-overlapping windows into archaic ancestry and non-archaic ancestry depending on the derived allele density.

#### Outgroup variants set, window mutation rate and callability and derived allele polarization

To generate the set of variants in the outgroup, we merged all variants from the following populations:

1. All Sub-Saharan Africans (populations: YRI, MSL, ESN) from the 1000 Genomes Project ^30^ and
2. All Sub-Saharan African populations from SGDP (this excludes Sharawi and Mozabite populations from the African supergroup) ^3^ except individuals from the Masai and Somali populations because they are reported to have some West Eurasian genetic component.

We determine the background mutation rate as the SNP density in the outgroup samples in windows of 100 kb.

To generate the callability regions, we merged the following files:

1. 1000 Genomes Project Callability file (hg19) ftp://ftp.1000genomes.ebi.ac.uk/vol1/ftp/release/20130502/supporting/accessible_genome_masks/StrictMask/
2. Repeatmask file (hg19) hgdownload.cse.ucsc.edu/goldenpath/hg19/bigZips/chromFaMasked.tar.gz To polarize alleles into ancestral and derived alleles we used the following file: http://web.corral.tacc.utexas.edu/WGSAdownload/resources/human_ancestor_GRCh37_e71/

#### Training the Hidden Markov model and decoding archaic fragments in each sample

For each extant non-African individual and 3 ancient samples (Stuttgart, Loschbour and Ust-ishim) from the SGDP, we filtered out all sites where the derived variant is found in our outgroup population and sites that are not in our callable regions.

Then we trained the HMM and found the best fitting emission and transition values. Finally we identified tracks of archaic introgression in the whole genome of each individual (Data1_archaicfragments.txt). The archaic fragments in Stuttgart, Loschbour and Ust-ishim are visualized in Fig. S1.

**Fig S1.**
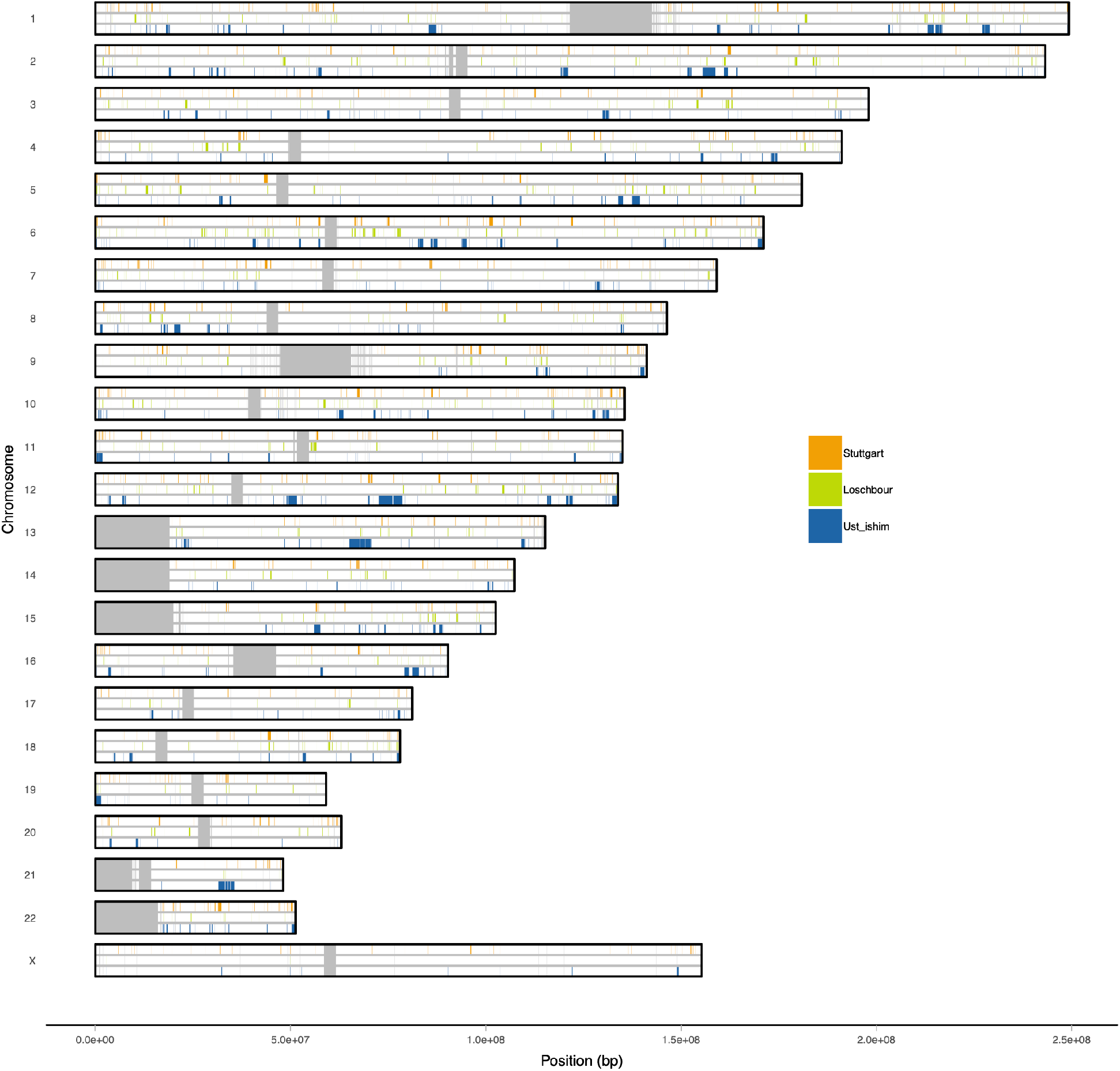
Archaic fragments in Ust *‘*Ishim, Loschbour and Stuttgart ancient samples. Each horizontal rectangle represents a chromosome (hg19). In each chromosome, it is shown the archaic fragments found in Ust *‘*Ishim, Loschbour and Stuttgart ancient samples (colour coded). Wide grey bands on the chromosomes show the non-callable portions of the genome (hg19).

### 4 Archaic fragment length gradient around the world is consistent to multiple filters

We studied the robustness of the difference in mean archaic fragment length among the 5 geographical groups studied applying multiple filters.

1. Median instead of mean The mean is very sensitive to outliers. In our case, very long archaic fragments, for example in East Asians, could increase the mean and thus show an unrealistic pattern among regions. To avoid that, we use median instead because it is more robust to outliers.
2. Vindija genome-like fragments The method used in this study is able to find archaic fragments whose variation is not fully captured by the sequenced archaic individuals ^11^. The difference in archaic fragment length can potentially be affected if there is a distinct archaic content among the extant populations studied here -for example, a greater and more recent Denisova component in Asia ^31^. It is known that the majority of the archaic component in Eurasia and America is from a Neanderthal population closely related to the Vindija genome ^12^. Thus, we restrict fragments used in this analysis to share more variation with the Vindija Neanderthal genome than the Altai Neanderthal genome or the Denisovan genome.
3. High confidence archaic fragments The method used in this study, returns the archaic fragments found in a genome with an associated mean posterior probability. We restricted archaic fragments compared to be of a high confidence (mean posterior probability >= 0.9). When we study the archaic fragment difference among individuals in Eurasia and America applying the multiple filters explained above, we can see that the pattern observed using all fragments holds (Extended Figure 1). We conclude that the difference in archaic fragment length is genuine and not depending on the factors exposed above.

### 5 Archaic fragment summary statistics per individual per region in extant populations and ancient samples

**Table S1.** **Archaic fragment summary statistics per individual per region**. Summary statistics of the fragments found among the individuals of the 5 main regions. For each statistic, the mean and the of SE (<u>S1</u>) is provided.

| Region | Number of samples | Number archaic fragments |  | Archaic seq (bp) |  | Mean archaic fragment length (bp) |  |
| --- | --- | --- | --- | --- | --- | --- | --- |
|  |  | mean | SE | mean | SE | mean | SE |
| West Eurasia | 71 | 980.92 | 6.93 | 72,134,504.74 | 747,098.22 | 73,449.52 | 375.37 |
| South Asia | 39 | 1,122.99 | 11.29 | 84,561,986.37 | 1,127,788.19 | 75,221.52 | 410.92 |
| America | 20 | 1,078.87 | 10.25 | 86,322,668.28 | 803,716.12 | 80,057.38 | 561.94 |
| Central Asia Siberia | 27 | 1,133.28 | 8.86 | 92,424,440.59 | 946,306.77 | 81,548.03 | 460.74 |
| EastAsia | 45 | 1,161.58 | 6.65 | 95,552,691.46 | 712,217.98 | 82,258.79 | 402.10 |

**Table S2.** **Archaic fragment summary statistics per ancient sample**. Summary statistics of the fragments found in the three ancient samples. For the archaic fragment length, the mean and the of SE (<u>S1</u>) is provided.

| Ancient samples | Number archaic fragments | Archaic seq (bp) | Archaic fragment length (bp) |  |
| --- | --- | --- | --- | --- |
|  |  |  | mean | SE |
| Ust 'Ishim | 763 | 134,360,000 | 176,100.09 | 12,464.19 |
| Loschbour | 921 | 75,757,000 | 82,190.02 | 3,087.97 |
| Stuttgart | 1,101 | 85,950,000 | 78,070.12 | 2,705.90 |

### 6 West Eurasia and East Asia fragment comparison

In this study, we compare fragments in West Eurasians and East Asians. The more individuals used to recover archaic fragments, the more undiscovered fragments can be found ^9^. Thus, the imbalance in the number of individuals in each region in the SGDP data (71 West Eurasians and 45 East Asians) can potentially affect any comparison between the two regions. Therefore, we downsample the number of individuals used in West Eurasians to 45 randomly chosen individuals to make comparisons fair.

First, we join all overlapping fragments for each region, hereby *“*joined region fragments*”* (Extended Figure 2). To do that, we used bedtools software ^32^ with the following command:

`bedtools merge -i ind1_regx.bed ind2_regx.bed … indN_regx.bed > joined_regx.bed`

where x denotes either West Eurasia or East Asia regions and N denotes the number of individuals in the corresponding region.

Then, we compared how much archaic sequence the two regions share (Extended Figure 2). For that, we call the intercept between the two joined sets of fragments. We refer to it as the *“*shared joined region sequence*”*. We use the following command:

`bedtools intercept -a joined_regx.bed -b joined_regy.bed > shared_joined.bed`

where x denotes either West Eurasia or East Asia and y denotes the other region different than x.

It follows that the rest of the fragments not included in this set are the *“*private joined region sequence*”*.

The amount of sequence for shared, private and total joined region fragments are provided in Table S3.

For each individual, we classified the fragments as shared depending upon if there was an overlapping fragment in the other joined region fragments (Extended Figure 2). We name these fragments as *“*shared individual fragments*”*. To get them, we ran the following command:

`beedtools intercept -u -a indn_regx.bed -b joined_regy.bed > shared_indn_regx.bed`

It follows that the rest of the fragments not included in this set are the *“*private individual fragments*”*.

Summary statistics for shared and private individual fragments are provided in Table S4.

Finally, we calculated the number of individuals that have an overlapping archaic fragment in a certain 1kb window in the genome. This way, we calculate the **archaic frequency**. For that, we first divided each fragment in the joined region fragments into 1 kb segments (joined_regx_1kb.bed). Then, we counted the number of individuals with an overlapping archaic fragment for each 1kb segment with the following command:

`bedtools intersect -c -a joined_regx_1kb.bed -b ind1_regx.bed ind2_regx.bed … indN_regx.bed > freq_regx.bed`

Extended Figure 3 shows a summary of the joined region fragments, shared joined region sequence and the archaic frequency for each region.

#### Shared joined region fragments filtering by archaic affinity

The collapsed East Asian archaic sequence (916,369,000 bp) is 1,06 fold greater than the collapsed West Eurasian archaic sequence (866,945,000 bp) and more than half of the sequence is shared between the two (485,255,000 bp, Table S5). We partially attribute this difference to the fact that East Asians have a higher Denisova component than West Eurasians ^31^. To study that we repeated the analysis above filtering archaic fragments in each individual (before collapsing) depending on which of the three archaic genomes (Vindija Neanderthal genome ^12^, Altai Neanderthal genome ^33^, Denisova genome ^34^) share the most variants to (below), following the methods in ^9^. Some fragments do not share variants with any of the 3 sequenced archaic genomes, and thus we classify them as unknown. There are also instances in which an archaic fragment does not share more SNPs with one of the archaic genomes but multiple, so we can*’*t classify the affinity of the fragments; these fragments are called ambiguous fragments.

1. Denisova fragments We only include archaic fragments which share more variants to Denisova genome than any of the two Neanderthal genomes.
2. nonDenisova fragments In this analysis we exclude fragments used above from all the fragments. Thus, we include Vindija-like, Altai-like, ambiguous and unknown.
3. Neanderthal fragments We only include archaic fragments that share more variants with either the Altai Neanderthal or the Vindija Neanderthal genomes than the Denisova genome. Neanderthal ambiguous fragments, fragments that share the same number of SNPs with Vindija or Altai but this number is higher than what is shared with the Denisova, are also included.

All results for the different filters are shown in Table S5. The Denisova content is 3 times greater in East Asia than in West Eurasia (Denisova fragments filter). When this unequal component is removed (non-Denisova fragments filter), we can see that the collapsed archaic sequence is very similar between the two regions.

The analysis was repeated with fragments that share more variation with Neanderthal than with Denisova (Neanderthal fragments). In this case, we observe a 1.07 fold higher Neanderthal content in the East Asian group. We attribute this to the fact that since West Eurasia archaic fragments tend to be shorter, they do not contain enough SNPs to classify them to the category that they belong to. Thus, they are going to be more often classified as unknown compared to fragments in East Asia. Furthermore, the ^11^ method has higher false negative rate with short fragments, which will artificially decrease the total number of fragments in that region.

**Table S3.** Summary table of shared, private and total joined archaic sequence of West Eurasia and EastAsia regions. Percent in respect of the total are shown in parenthesis.

| Region | Number of samples | Type | Archaic Sequence (kb) |
| --- | --- | --- | --- |
| West Eurasia | 45 | Shared | 485,255<br>(55,97%) |
|  |  | Private | 381,690<br>(44,03%) |
|  |  | All | 866,945<br>(100%) |
| East Asia | 45 | Shared | 485,255<br>(52,95%) |
|  |  | Private | 431,114<br>(47,05%) |
|  |  | All | 916,369<br>(100%) |

**Table S4.** Summary statistics of the shared, private and total individual archaic fragments of West Eurasians and East Asians. For each statistic, the mean and the of SE (<u>S1</u>) is provided.

| Region | Number of samples | Type | Number archaic fragments |  | Archaic seq (bp) |  | Archaic fragment length (bp) |  |
| --- | --- | --- | --- | --- | --- | --- | --- | --- |
|  |  |  | mean | SE | mean | SE | mean | SE |
| West Eurasia | 45 | Shared | 756.17 | 9.43 | 59,867,254.64 | 945,255.36 | 79,060.32 | 473.90 |
|  |  | Private | 221.48 | 2.83 | 11,477,725.85 | 239,923.71 | 51,722.32 | 737.11 |
|  |  | All | 977.85 | 9.83 | 71,357,270.60 | 988,585.16 | 72,879.08 | 447.28 |
| East Asia | 45 | Shared | 913.80 | 5.04 | 81,720,490.56 | 586,860.89 | 89,448.78 | 473.32 |
|  |  | Private | 247.80 | 3.72 | 13,824,416.74 | 249,255.72 | 55,785.00 | 573.58 |
|  |  | All | 1161.57 | 6.76 | 95,555,705.32 | 704,807.56 | 82,258.99 | 400.50 |

**Table S5.** Joined archaic sequence in East Asia and West Eurasia and comparative statistics for different subsamples of archaic fragments (<u>S6</u>).

|  | Joined East Asia archaic sequence (kb) | Joined West Eurasia archaic sequence (kp) | Fold diff | Shared joined archaic sequence (kb) | East Asia shared (%) | West Eurasia shared (%) |
| --- | --- | --- | --- | --- | --- | --- |
| All fragments | 916,369 | 866,945 | 1.06 | 485,255 | 52.95 | 55.97 |
| Denisova fragments | 107,695 | 36,850 | 2.92 | 16,004 | 14.86 | 43.43 |
| nonDenisova fragments | 853,065 | 850,028 | 1.003 | 460,490 | 53.98 | 54.17 |
| Neanderthal fragments | 646,710 | 604,518 | 1.07 | 309,043 | 47.79 | 51.12 |

### 7 Derived alleles call outside regions with evidence of archaic introgression and acquired after the Out of Africa in SGDP samples

We retrieved the genotypes of all polymorphic loci for each individual in the 5 main regions and African samples using the cpoly script from the Ctools software^3^ for chromosomes 1 -22. In the parameter file, we specified the minimum quality to be 1 (as recommended by^3^) and alleles to be polarized with the chimpanzee reference genome (PanTro2) provided with the SGDP data.

Next, we masked repetitive regions and regions of the genome in which there is some evidence of archaic introgression.

1. Neandertal introgressed regions Neanderthals had a different mutation profile than modern humans^9^. Thus, differences in Neanderthal content per individual could influence those analyses that explore the mutation spectrum differences among populations. Also, by removing these regions, we will base the mutation analysis on regions of the genome that we haven*’*t explored in the archaic fragment length part of the study. Thus, the tests are going to be independent of each other. To do that, we disregarded any polymorphism localized in a region with evidence of archaic introgression in any of the individuals analyzed in this study (S3). For that, we joined all archaic fragments called in any individual included in this study using this command: `bedtools merge -i ind1.bed ind2.bed … indN.bed > joined.bed` where N denotes the total number of individuals. In total, the joined archaic region adds up to 1,632,776,000 bp.
2. Repeats We also excluded repetitive regions in which sequencing errors are expected to be more prevalent. For that, we downloaded the human reference genome by using the following command: `for chr in ‘seq 1 22’ X Y;` `do` `rsync -avzP` `rsync://hgdownload.cse.ucsc.edu/goldenPath/hg19/chromosomes/chr${chr}.fa.gz .; done`

from which we created a bed file with the coordinates of the repeats from RepeatMasker and Tandem Repeats Finder (represented in the reference genomes fastas as lowercase letters in the fasta file).

These regions add up to 1,431,504,380 bp in total.

The intersection between the repetitive regions and the archaic regions correspond to 806,042,777 bp, which corresponds to 56.31% of the total repetitive regions sequence and 49.37% of the archaic sequence. Together, these regions add up to 2,258,237,603 bp. If we consider only the callable fraction -instead of the total genomic length of 3,036,303,846 bp -of the human genome (2,835,673,565 bp), 577,435,962 bp remain after masking by archaic and repetitive regions (20.36%).

Other filters on the SNP level were imposed for each polymorphism:

1. The SNP must be biallelic
2. The contiguous 5*’* and 3*’* base pairs of the focal SNP (context) must be called in the human reference genome (hg19)
3. 20% of the individuals have to be called
4. The chimpanzee reference genome in human coordinates must have the homologous base pair called for that position
5. No Sub Saharan African (which excludes S_Mozabite-1, S_Mozabite-2, S_Saharawi-1 and S_Saharawi-2 samples from the African supergroup) samples can have the derived allele

The latter filter ensures that the polymorphisms investigated most probably arose after the Out of Africa expansion. S_Masai-1, S_Masai-2 and S_Somali-1 samples are not included in the Sub Saharan African group because they are reported to have some West Eurasian genetic component in ^3^, which would affect our results. If African genomes with West Eurasian components are included in the African set, then, by the 5) filter, we are going to more likely remove derived alleles private to West Eurasia than other regions.

Homozygous locus for the derived allele count as 2 mutations and individuals heterozygous count as 1 for a given individual. The distribution of derived allele accumulation per region is shown in Fig. 3 and the mean derived allele accumulation counts per region are provided in Table S6.

**Fig S2.**
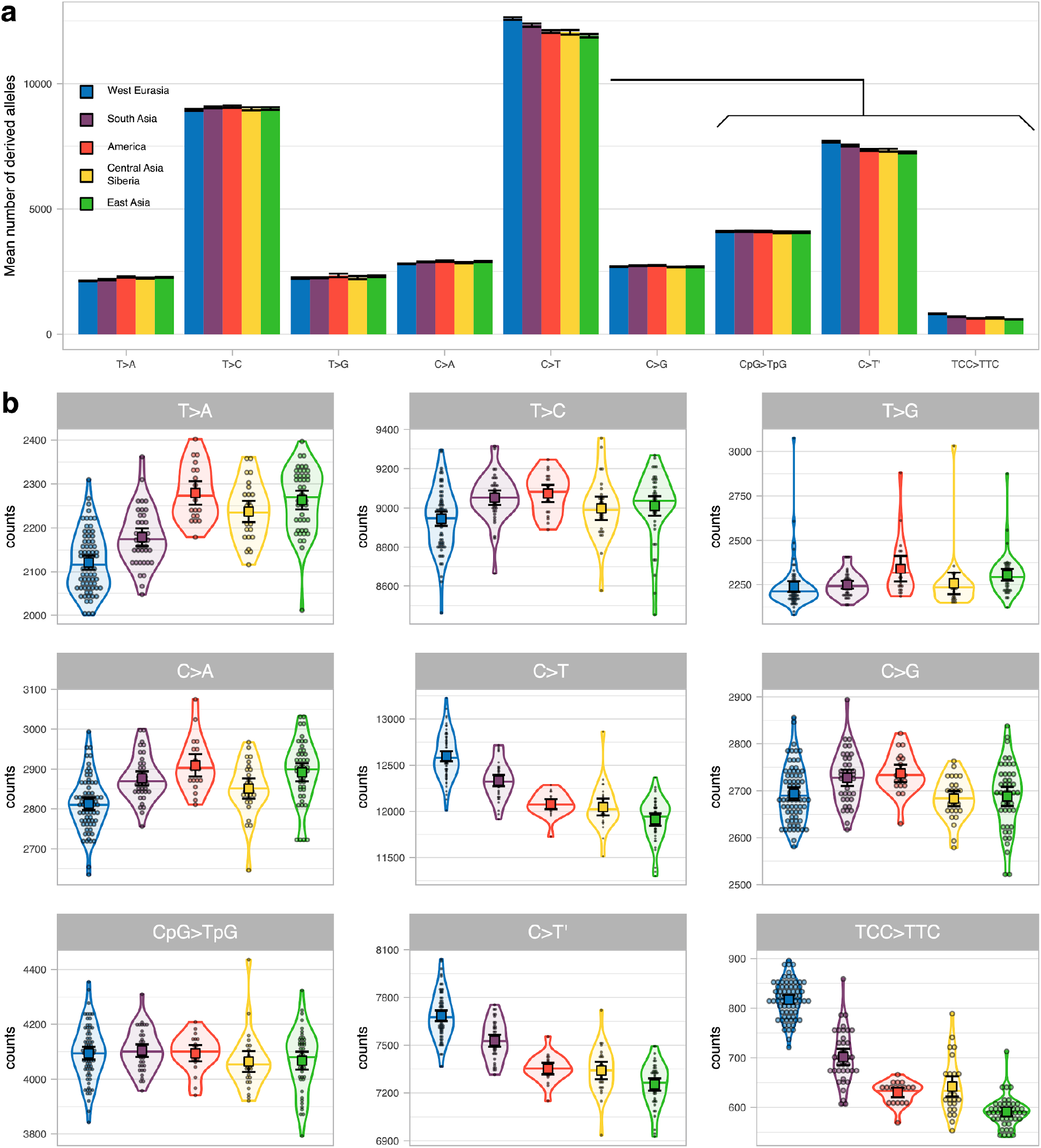
Mean derived allele accumulation of the 9-mutation types per region. **a)** The mean number of derived alleles of each mutation type accumulated among individuals of the 5 regions (colour coded). The 95%CI of each mean is shown as error bars. **b)** The number of derived alleles of each mutation type per region (colour coded) as violin plot. Individual values are shown as dots. The median is shown as a horizontal line in each violin plot. The mean and its 95%CI of each distribution is shown as a coloured square with their corresponding error bars.

**Fig S3.**
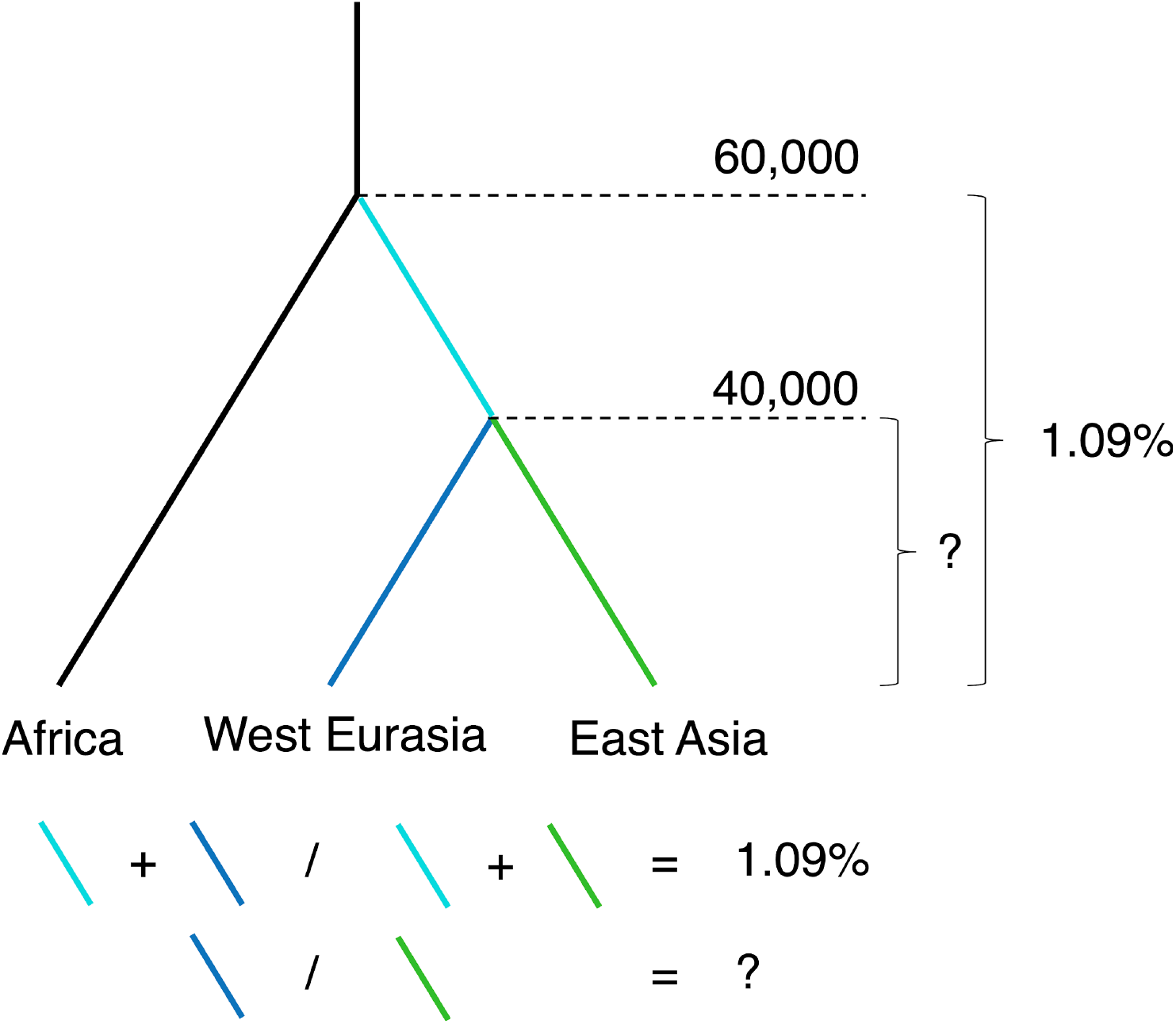
Mutation rate difference between West Eurasia and East Asia. This diagram shows conceptually that the mutation rate could only be different after the split between East Asians and West Eurasians (blue and green terminal branches). However, the difference in derived allele accumulation is calculated since the split with Africans for each group (cyan and blue, cyan and green).

**Table S6.** **Derived allele accumulation per region**. Summary statistics of the derived allele accumulation per region (<u>S7</u>). For each region, the mean and the of SE (<u>S1</u>) is provided.

| Region | Number of samples | Derived allele accumulation |  |
| --- | --- | --- | --- |
|  |  | mean | SE |
| West Eurasia | 71 | 31,408.54 | 62.61 |
| South Asia | 39 | 31,418.28 | 56.07 |
| America | 20 | 31,417.64 | 83.31 |
| Central Asia Siberia | 27 | 31,074.21 | 88.53 |
| EastAsia | 45 | 31,069.77 | 80.99 |

**Table S7.** **Derived allele accumulation per region stratified per mutation type**. Summary statistics of the derived allele accumulation per region for each mutation type (<u>S7</u>). For each region and mutation type, the mean and the of SE (<u>S1</u>) is provided.

| Region | T |  |  |  |  |  |
| --- | --- | --- | --- | --- | --- | --- |
|  | T>A |  | T>C |  | T>G |  |
|  | mean | SE | mean | SE | mean | SE |
| West Eurasia | 2,121.02 | 8.11 | 8,944.94 | 18.57 | 2,238.91 | 15.64 |
| South Asia | 2,178.71 | 10.37 | 9,052.32 | 18.91 | 2,249.05 | 10.88 |
| America | 2,279.45 | 13.67 | 9,073.85 | 22.37 | 2,340.34 | 37.02 |
| Central Asia<br>Siberia | 2,237.11 | 12.33 | 8,997.90 | 30.43 | 2,257.22 | 31.10 |
| EastAsia | 2,263.41 | 10.86 | 9,009.86 | 25.46 | 2,305.85 | 17.52 |

| Region | C |  |  |  |  |  |
| --- | --- | --- | --- | --- | --- | --- |
|  | C>A |  | C>T |  | C>G |  |
|  | mean | SE | mean | SE | mean | SE |
| West Eurasia | 2,812.88 | 7.93 | 12,597.01 | 27.81 | 2,693.51 | 6.96 |
| South Asia | 2,876.76 | 8.80 | 12,333.87 | 31.10 | 2,727.42 | 8.86 |
| America | 2,909.30 | 14.36 | 12,077.68 | 27.39 | 2,737.04 | 9.47 |
| Central Asia<br>Siberia | 2,851.11 | 13.00 | 12,047.72 | 46.23 | 2,683.31 | 8.42 |
| EastAsia | 2,891.90 | 11.58 | 11,911.38 | 34.26 | 2,687.80 | 10.43 |
|  | CpG>TpG |  | T>C' |  | TCC>TTC |  |
|  | mean | SE | mean | SE | mean | SE |
| West Eurasia | 4,094.57 | 11.82 | 7,684.86 | 17.14 | 817.35 | 4.42 |
| South Asia | 4,103.80 | 11.75 | 7,528.27 | 19.06 | 701.81 | 8.40 |
| America | 4,094.27 | 14.91 | 7,353.29 | 18.45 | 629.95 | 4.81 |
| Central Asia<br>Siberia | 4,064.23 | 19.55 | 7,341.42 | 28.09 | 642.05 | 10.48 |
| EastAsia | 4,067.34 | 16.60 | 7,252.96 | 19.80 | 591.16 | 4.48 |

Finally, we classified loci in different mutation types depending on the derived allele nucleotide, the ancestral allele nucleotide and their 5*’* and 3*’* nucleotide context. For example, as shown by the diagram below, a derived allele T that had an ancestral allele C with the context G and A (5*’* and 3*’* respectively) would be denoted as GCA>T. Because we do not make distinction of the strand in which the mutation occurred, we collapsed strand-symmetric mutations. This is the same as saying that GCA>T is equivalent to TGC>A. This way, we end up with 96 mutation types.

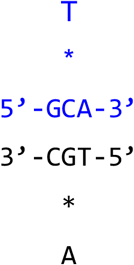

Data2_mutationspectrum.txt provides the resulting counts of each individual for each mutation type in each chromosome.

The mutation types investigated in this study are the 9:

- 6 mutation types in which only the ancestral and derived allele nucleotides were taken into account and C and T were used as ancestral (T>A, T>C, T>G, C>A, C>T, C>G)
- C>T mutations were further divided into 3 mutation types:
  - CpG>TpG mutations which are shown to evolve in a more clock like manner^24^.
  - TCC>TTC mutations which are in excess in Europeans compared to other human populations^5,22^.
  - C>T*’* mutations which contain the rest of C>T mutations not included in the previous 2 types.

The distribution of derived allele accumulation per region is shown in Fig. S2 and the mean derived allele accumulation counts per region are provided in Table S7.

### 8 Estimation of the different parental generation time in West Eurasia and East Asia

As described in the main text, West Eurasia individuals have accumulated 1.09% more derived alleles than East Asians since the split with Africans (Out of Africa). Because we are only interested in the proportion of derived alleles accumulated after the split of West Eurasians and East Asians, we need to correct for the span of time since the Out of Africa event until the split of the two Eurasian populations (Fig. S3). Thus, we need to assume dates for the split between Africans and non-Africans and the split between Eurasians.

We note that in the literature dating the Out of Africa is widely discussed and controversial, since it was not a clean split between non-Africans and Africans. Instead, from MCMC results and cros coalescence rate analysis in ^2,19^ the authors note that there might have been a gradual separation among African populations and between Africans and non-Africans. They suggest that this process created population structure between 200,000 -100,000 years ago within Africa and that the non-African group had more gene flow with certain African groups (i.e., Yorubans) than others (i.e., San). After that, the rate increased, indicative of an accelerated split between Africans and non-Africans which has the median divergence point between 80,000 -60,000 years ago. Similarly, the split among Eurasians was not clean either. All splits started around 70,000 years ago with a median divergence point between 40,000 and 20,000 years ago for East Asians and West Eurasians. Nonetheless, studies of ancient DNA show that around 40,000 years ago East Asians and West Eurasians were already diverging: the ancient human sample of Kostenki (36,000 year old sample) presents higher affinity to present day West Eurasians ^21^ and Tianyuan (40,000 year old sample) to East Asians ^20^.

In this analysis, we assume that the split between Africans and non-Africans happened 60,000 years ago and that the split between West Eurasians and East Asians happened 40,000 years ago. This is because if the proportion of time the West Eurasians and East Asians were apart decreases in respect of the time since both splited from Africans (i.e., out of Africa happening 80,000 instead of 60,000 years ago), the rate at which mutations should have accumulated would have been higher. Thus, a conservative measurement will be assuming a lower bound for the out of Africa.

In consequence, the excess of derived alleles accumulated in West Eurasians compared to East Asia is:

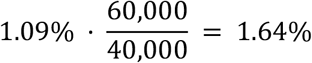

In ^4^ a poisson regression is derived for the number of mutations transmitted in each generation from trio data for each parental lineage depending on their age at reproduction:

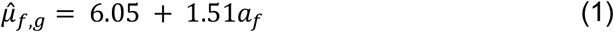

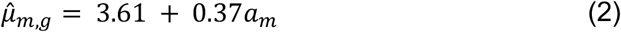

Where subscripts *f* and *m* denote paternal and maternal respectively, 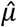 is the estimation of the mean mutation rate per generation (*g*) and *a* is the mean parental age. Thus, assuming the same mean parental age for both progenitors (*a*_*f*_ = *a*_*m*_ = *a*) we get that the total mutation rate per generation is calculated by the equation 3 and the yearly (*y*) rate by equation 4.

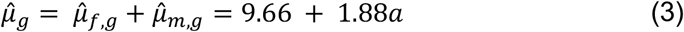

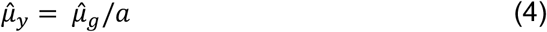

Then, to compare the mutation rate per year in two different populations (*x* and *z*) with different mean parental ages, we get that

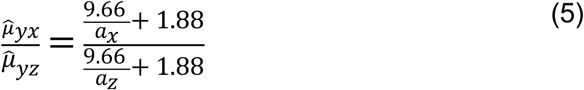

The number of derived alleles accumulated in a genome during a period of time (*d*) depends on the mutation rate per year and the time span (*T*)

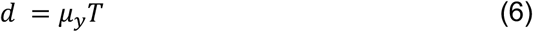

However, the ratio of *d* between two populations, will only depend on their mutation rate because *T* has been the same for both

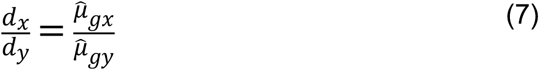

Thus, we can estimate the *a*_*x*_if *a*_*z*_and the *d*_*x*_*/d*_*z*_are known

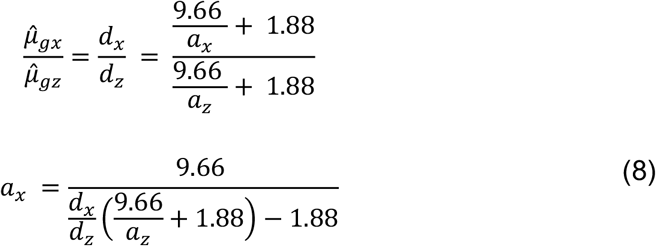

In this study, we find that the ratio of the mean derived allele accumulation in West Eurasia (*WE*) vs East Asia (*EA, d* _*WE*_ */d*_*EA*_) is 1.0164 (1.64%). With formula (8), we check for reasonable *a*_*EA*_ values between 28 and 32 years and found that the values of *a*_*WE*_ ranged between 25.32 and 28.59 respectively. Thus, we estimate that generation times in East Asians have been 2.68 to 3.39 years longer than in West Eurasians since the split of the two populations. This corresponds to West Eurasians having had approx. 150 generations more than East Asians.

**Fig S4.**
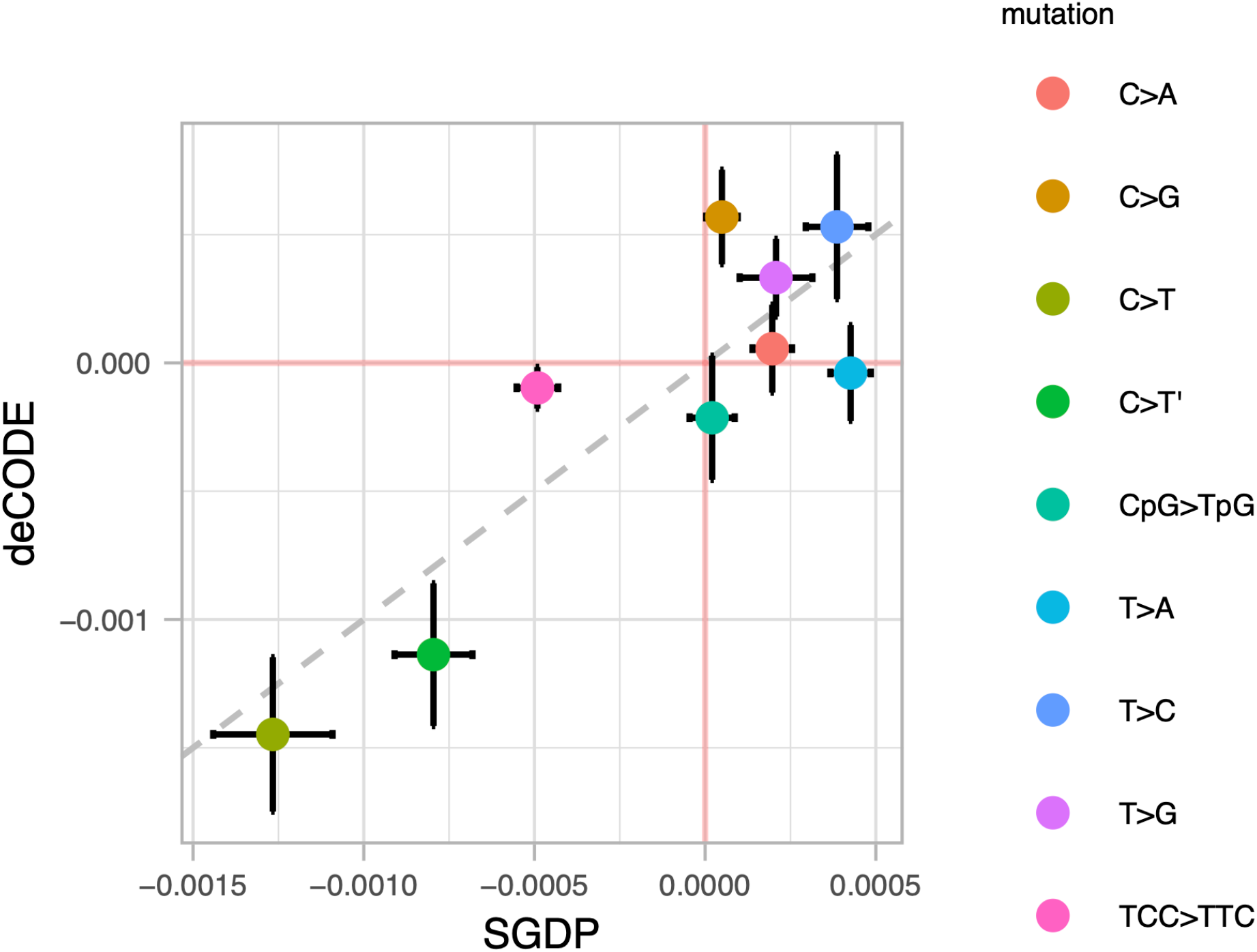
Slope coefficient correlation between SGDP data and deCODE data linear models. Dot plot graph illustrating the correlation between linear model slope coefficients derived from the SGDP data (x-axis) and deCODE data (y-axis) for each mutation type (color code). 95%CI for each estimate are shown as error bars. The 1-to-1 correspondence is denoted by the gray dashed diagonal line.

**Fig S5.**
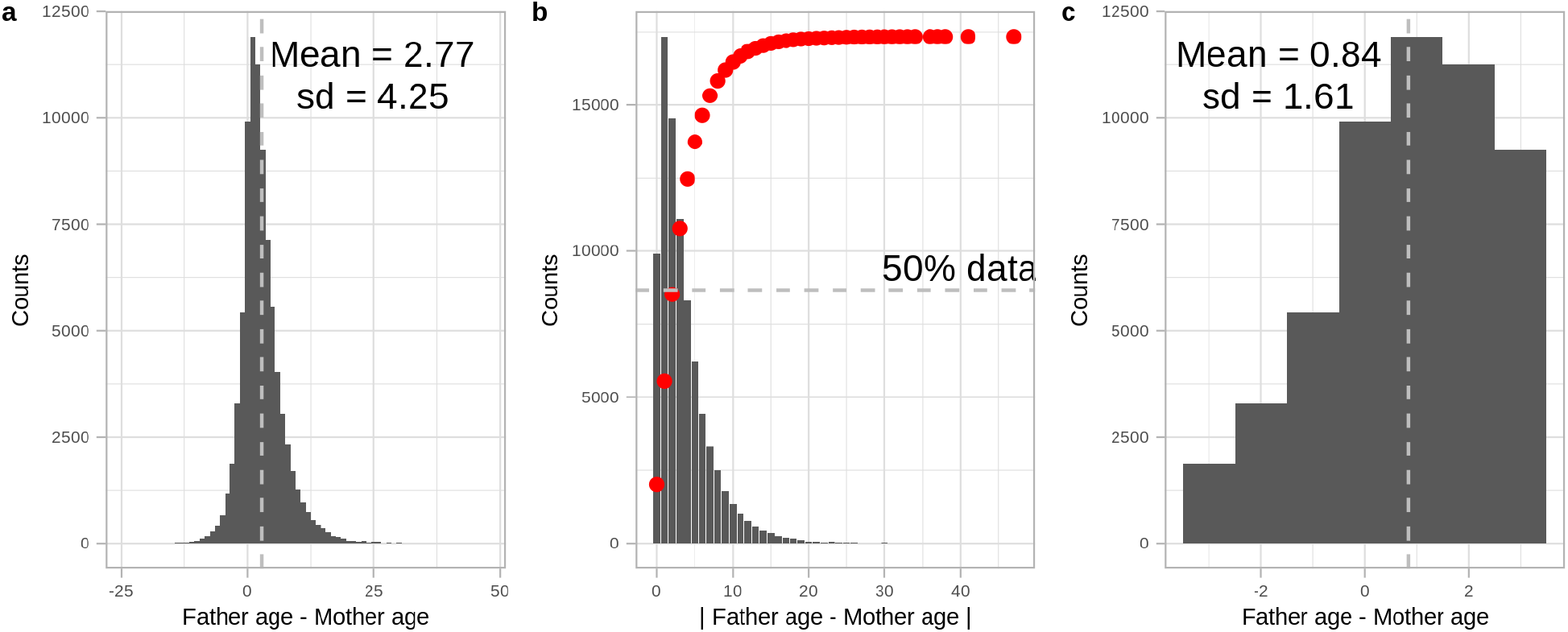
Parental age difference in the deCODE data. **a)** Histogram of the number of probands with a certain parental age difference. The mean is shown as a vertical gray line and annotated as a numeric figure. **b)** Histogram of the number of probands with a certain absolute parental age difference. The cumulative distribution of provands is denoted by red dots. The horizontal gray line shows the 50% data threshold. **c)** Histogram of the number of probands with a certain parental age difference with less than 4 years difference. The mean is shown as a vertical gray line and annotated as a numeric figure.

**Fig S6.**
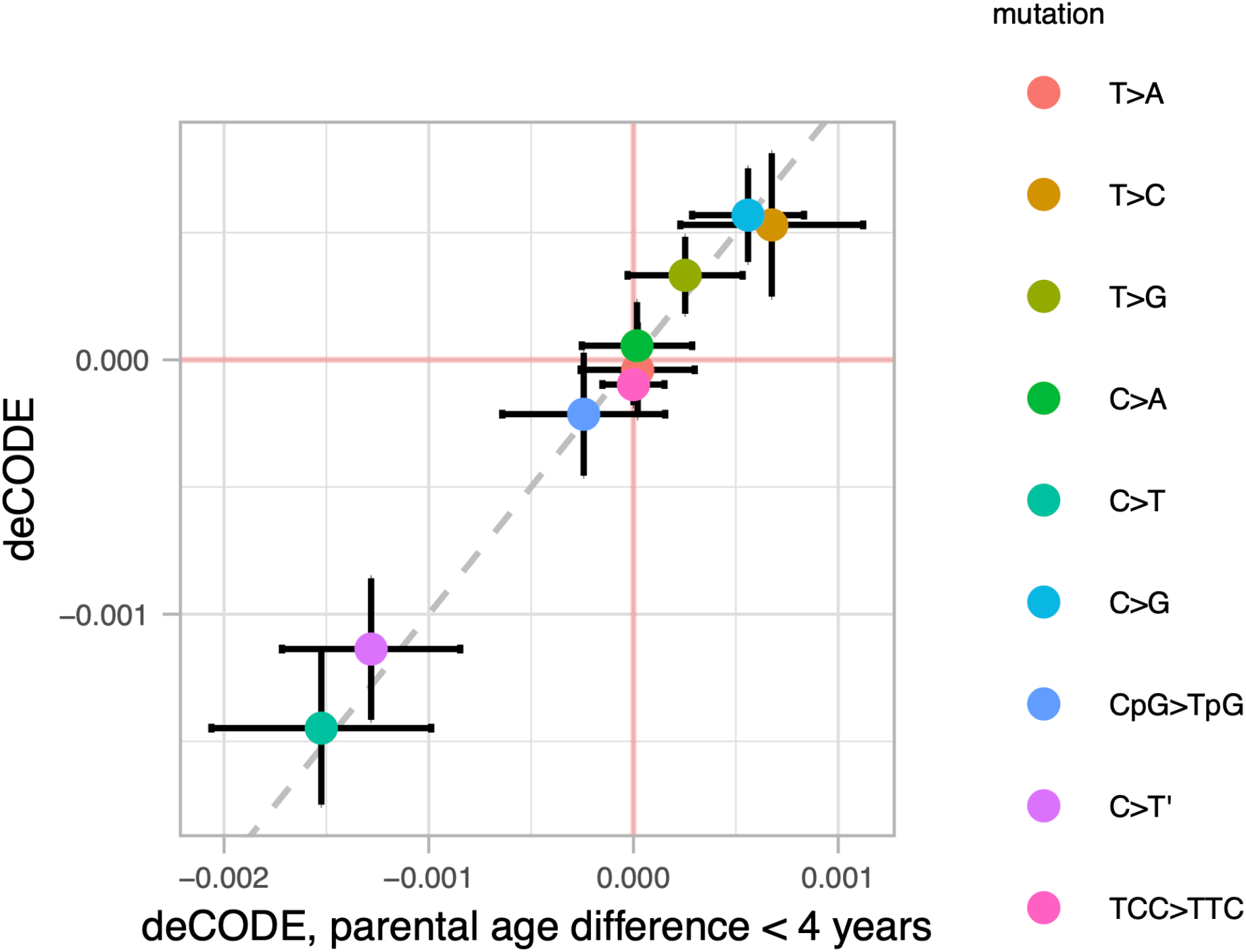
Slope coefficient correlation between linear models of deCODE data and deCODE data when only using probands with parental age difference less than 4 years. Dot plot graph illustrating the correlation between linear model slope coefficients derived from the deCODE data (y-axis) and the deCODE data when only using probands with parental age difference less than 4 (x-axis) for each mutation type (color code). 95%CI for each estimate are shown as error bars. The 1-to-1 correspondence is denoted by the gray dashed diagonal line.

**Table S8.**
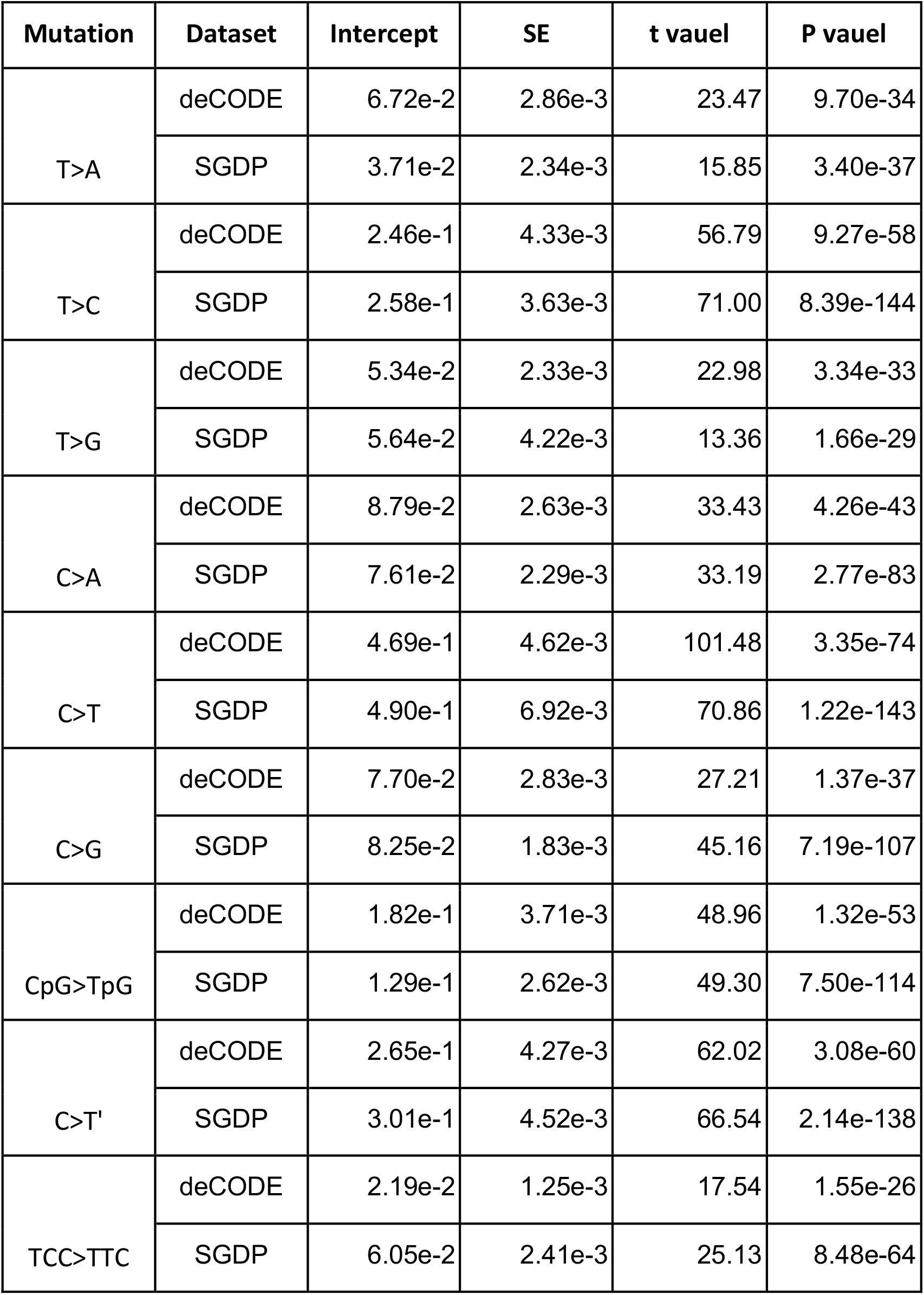
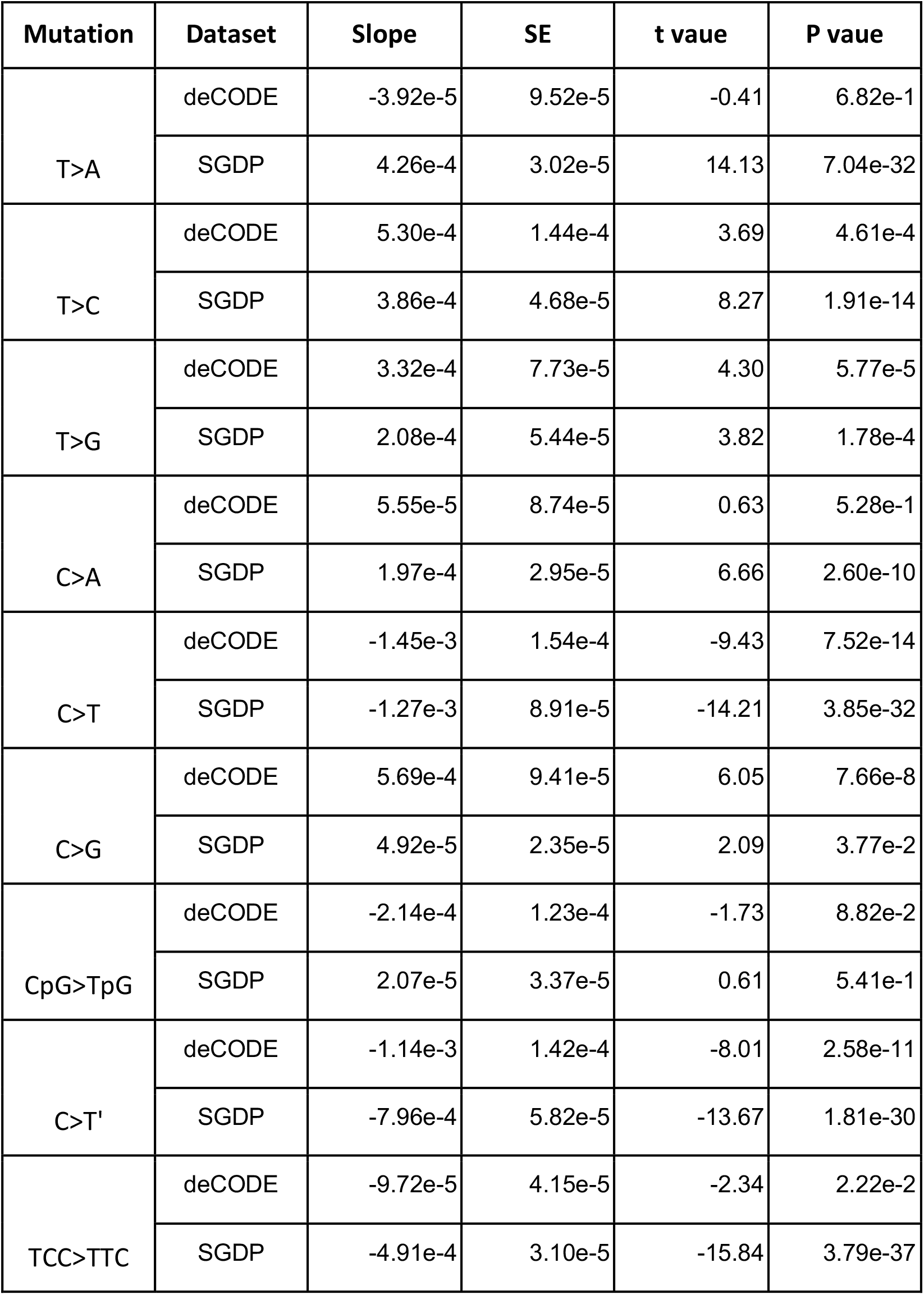
**Linear models between mutation type fraction and mean generation time estimate in the SGDP and deCODE data sets**. Two separate tables are given for the intercept and the slope of the linear models. For each mutation type and data set, the coefficients estimate the SE, t value and the associated P value are provided.

| Mutation | Dataset | Intercept | SE | t vauel | P vauel |
| --- | --- | --- | --- | --- | --- |
| T>A | deCODE | 6.72e-2 | 2.86e-3 | 23.47 | 9.70e-34 |
|  | SGDP | 3.71e-2 | 2.34e-3 | 15.85 | 3.40e-37 |
| T>C | deCODE | 2.46e-1 | 4.33e-3 | 56.79 | 9.27e-58 |
|  | SGDP | 2.58e-1 | 3.63e-3 | 71.00 | 8.39e-144 |
| T>G | deCODE | 5.34e-2 | 2.33e-3 | 22.98 | 3.34e-33 |
|  | SGDP | 5.64e-2 | 4.22e-3 | 13.36 | 1.66e-29 |
| C>A | deCODE | 8.79e-2 | 2.63e-3 | 33.43 | 4.26e-43 |
|  | SGDP | 7.61e-2 | 2.29e-3 | 33.19 | 2.77e-83 |
| C>T | deCODE | 4.69e-1 | 4.62e-3 | 101.48 | 3.35e-74 |
|  | SGDP | 4.90e-1 | 6.92e-3 | 70.86 | 1.22e-143 |
| C>G | deCODE | 7.70e-2 | 2.83e-3 | 27.21 | 1.37e-37 |
|  | SGDP | 8.25e-2 | 1.83e-3 | 45.16 | 7.19e-107 |
| CpG>TpG | deCODE | 1.82e-1 | 3.71e-3 | 48.96 | 1.32e-53 |
|  | SGDP | 1.29e-1 | 2.62e-3 | 49.30 | 7.50e-114 |
| C>T' | deCODE | 2.65e-1 | 4.27e-3 | 62.02 | 3.08e-60 |
|  | SGDP | 3.01e-1 | 4.52e-3 | 66.54 | 2.14e-138 |
| TCC>TTC | deCODE | 2.19e-2 | 1.25e-3 | 17.54 | 1.55e-26 |
|  | SGDP | 6.05e-2 | 2.41e-3 | 25.13 | 8.48e-64 |

**Table S8. Linear models between mutation type fraction and mean generation time estimate in the SGDP and deCODE data sets.**
| Mutation | Dataset | Slope | SE | t vaue | P vaue |
| --- | --- | --- | --- | --- | --- |
| T>A | deCODE | -3.92e-5 | 9.52e-5 | -0.41 | 6.82e-1 |
|  | SGDP | 4.26e-4 | 3.02e-5 | 14.13 | 7.04e-32 |
| T>C | deCODE | 5.30e-4 | 1.44e-4 | 3.69 | 4.61e-4 |
|  | SGDP | 3.86e-4 | 4.68e-5 | 8.27 | 1.91e-14 |
| T>G | deCODE | 3.32e-4 | 7.73e-5 | 4.30 | 5.77e-5 |
|  | SGDP | 2.08e-4 | 5.44e-5 | 3.82 | 1.78e-4 |
| C>A | deCODE | 5.55e-5 | 8.74e-5 | 0.63 | 5.28e-1 |
|  | SGDP | 1.97e-4 | 2.95e-5 | 6.66 | 2.60e-10 |
| C>T | deCODE | -1.45e-3 | 1.54e-4 | -9.43 | 7.52e-14 |
|  | SGDP | -1.27e-3 | 8.91e-5 | -14.21 | 3.85e-32 |
| C>G | deCODE | 5.69e-4 | 9.41e-5 | 6.05 | 7.66e-8 |
|  | SGDP | 4.92e-5 | 2.35e-5 | 2.09 | 3.77e-2 |
| CpG>TpG | deCODE | -2.14e-4 | 1.23e-4 | -1.73 | 8.82e-2 |
|  | SGDP | 2.07e-5 | 3.37e-5 | 0.61 | 5.41e-1 |
| C>T' | deCODE | -1.14e-3 | 1.42e-4 | -8.01 | 2.58e-11 |
|  | SGDP | -7.96e-4 | 5.82e-5 | -13.67 | 1.81e-30 |
| TCC>TTC | deCODE | -9.72e-5 | 4.15e-5 | -2.34 | 2.22e-2 |
|  | SGDP | -4.91e-4 | 3.10e-5 | -15.84 | 3.79e-37 |

### 9 Mutation spectrum correlation with mean parental age

The germline mutation spectrum is dependent on the parental sex and age at conception ^4^. In this study, we observe differences in the abundance of derived alleles accumulated after the out of Africa event when stratified by mutation type (Fig. S2, Table S7). Here, we study to which extent these differences can be explained by changes in generation time in the 5 regions. For that, we compare the mutational patterns of *de novo* mutations (DNM) depending on parental age in trio studies ^4,25^ (deCODE dataset) with the differences in mutation spectrum of extant populations with the mean archaic fragment length as a proxy of mean generation time (SGDP dataset).

#### SGDP dataset

We classified the derived alleles found in the autosomes of each individual into 6 mutation types depending on the ancestral and derived allele as explained in S7. C>T mutations were also classified in 3 subtypes: TCC>TTC, CpG>TpG and the rest (C>T*’*). In total, we divide all mutations into 9 types. In order to obtain the fraction of each mutation type per individual, we divided the number of each mutation type by the total amount of derived alleles. C>T mutations are duplicated since we subdivide them into 3 extra categories (TCC>TTC, CpG>TpG and C>T*’*). Thus, the total amount of derived alleles do not consider these 3 types. We correlated the fraction of derived alleles of each type with the mean archaic fragment length as a proxy of mean generation time (Fig. 3b). We obtained the linear model of such correlation for each mutation type using the following R function (Table S8).

`lm(mutation_fraction∼mean_fragment_length)`

#### deCODE dataset

We downloaded the set of DNM called in ^25^ and the additional proband information from the supplementary data provided in the publication. We join both in order to compute the mean parental age for each DNM for each proband. Indels are filtered out. Following the methodology in a similar test in ^4^, we aggregate all mutation counts for each of the 9 types of all probands with the same mean parental age. We then compute the fraction of each mutation type. In other words, for each mutation type and mean parental age we have a single mutation fraction value. Those data points that were obtained aggregating information from less than 2 probands were discarded. We obtained linear models for each mutation type using the following R function (Table S8).

`lm(mutation_fraction∼mean_parental_age, weights = n_probands)`

##### The correlations between the slopes of both datasets is shown in Fig. S4

The probands of the deCODE dataset have a bias towards fathers being older than mothers, with a mean of 2.77 years and the largest difference of more than 40 years (Fig. S5a). To study if the correlation of mutation spectrum with the mean parental age is affected by the mentioned bias, we rerun the correlation test with the deCODE dataset with only probands that have parents with an age difference of less than 4 years. This way, we retaining more than 50% of the data (Fig. S5b) and reduce the bias (mean = 0.94 differences in years, Fig. S5c). We then compared the slopes of the linear models calculated in the original deCODE dataset and when we impose the parental age difference filter explained above (Fig. S6). We don*’*t observe qualitative changes in the slopes when comparing the two and thus, we used all probands for our analysis.

## 10 Sex Specific mutational patterns

### X-to-A ratio

Due to the inheritance pattern of the X chromosome -2 copies transmitted in females while only 1 in males -compared to autosomes -2 copies in both females and males -, it is expected that the X chromosome has ¾ the diversity of the autosomes. However, this can be altered if the mutation rate changes disproportionately between females and males due to shifts in generation time between sexes. For example, an increase in the male mean generation time will decrease the yearly mutation rate in males and thus, proportionally less mutations are going to be accumulated in autosomes compared the X chromosomes ^28^. Therefore, the ratio of derived allele accumulation between the X chromosome and the autosomes will reflect variation on the generation time between males and females: higher values of the X-to-A ratio will be indicative of longer generation times in males compared to females and vice versa. Although here we only consider generation time differences to affect the ratio, there are other factors that can perturbe this ratio such as reproductive variance between sexes ^35^, demographic events ^36^ or differences in selection ^37^.

To investigate that, we obtained the number of derived alleles in the autosomes and X chromosomes of the females of the SGDP data (Table S9), as described in S7 (included in Data2_mutationspectrum.txt), and computed the X-to-A ratio as:

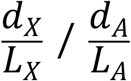

where *d* denotes the number of derived alleles, *L* the number of callable base pairs in either *X*(X chromosome) or *A*(autosomes). We then correlated the ratio with the mean archaic fragment length for each individual obtained in S3 (Fig. 4a).

### C>G maternaly enriched regions

As described in ^4^, there are regions of the genome in which DNM are clustered (cDNM). Those regions appear to be enriched in C>G mutations which originate in the maternal lineage. They also show that these clusters increase in number more rapidly with maternal than paternal age at conception.

Here we explore if there is a difference on the number of C>G segregating sites in cDNM genomic windows among the 5 regions.

For that we compute the number of derived alleles that are C>G and non-C>G along the genome in windows of 1Mb. We join this information with the annotation of 1Mb-window of the genome as cDNM or non-cDNM provided in ^4^. Then, for each individual we compute the following:

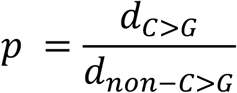

where *d* denotes the number of derived alleles of C>G or non-C>G. Thus, p is the ratio between the two quantities. Then, to compare this ratio between cDNM and non-cDNM regions we compute the mean *p* (*p*) over all regions and compute the following ratio

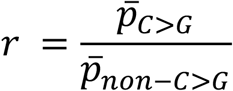

If *r* = 1, it shows that there are a similar number of C>G mutations in cDNM regions compared to the rest of the genome. If *r* > 1, then there is an excess and if *r* < 1, then there is a depletion. Nonetheless, we are not interested in the actual ratio, but the comparison among regions on this quantity. We then correlated the ratio with the mean archaic fragment length for each individual obtained in S3 (Fig. 4b).

### Y chromosome

Male individuals with shorter generation time are predicted to increase the mutation rate per year. Thus, Y chromosomes are expected to accumulate more derived alleles in individuals with a historically shorter mean generation time compared to others with longer ones.

To investigate that, we followed a similar procedure as in S7, changing certain steps and filters listed below:

1. We only used males in SGDP data
2. Alleles were polarized using the Chimp sequence in human coordinates. Since the chimpanzee Y chromosome is not provided with the SGDP data, this was achieved by taking the chimpanzee sequence from the hg19-panTro6 alignment into a fasta file with the human coordinates. The alignment can be downloaded from the following link: http://hgdownload.cse.ucsc.edu/goldenpath/hg19/vsPanTro6/reciprocalBest/axtRBestNet/hg19.panTro6.rbest.axt
3. No archaic regions were masked since there is no evidence of archaic sequence in the modern human Y chromosome
4. Only polymorphisms in the X degenerate regions are considered (coordinates from ^38^) and no further filters regarding repetitive regions were imposed
5. Individuals S_Finnish-2, S_Finnish-3, S_Palestinian-2, S_Mansi-1 and S_Masai-2 were discarded from the analysis because they didn’t yield any callable polymorphism
6. For each individual, all heterozygous sites were classified as non callable sites
7. Only African individuals with Y haplogroups A and B (metadata provided in ^3^, A: S_Ju_hoan_North-2, S_Dinka-2; B: S_Biaka-1, S_Biaka-2, S_Mbuti-3, S_Ju_hoan_North-3, S_Ju_hoan_North-1) were used as the outgroup. If polymorphisms were found to be segregating in these individuals, they were filtered out from this analysis
8. We didn*’*t require the 5*’* and 3*’* contiguous base pairs (context) of a polymorphic site to be callable

The accumulation of derived alleles in the Y chromosome per geographical region is shown in Fig. S7 (included in Data2_mutationspectrum.txt) and in Table S10.

**Fig S7.**
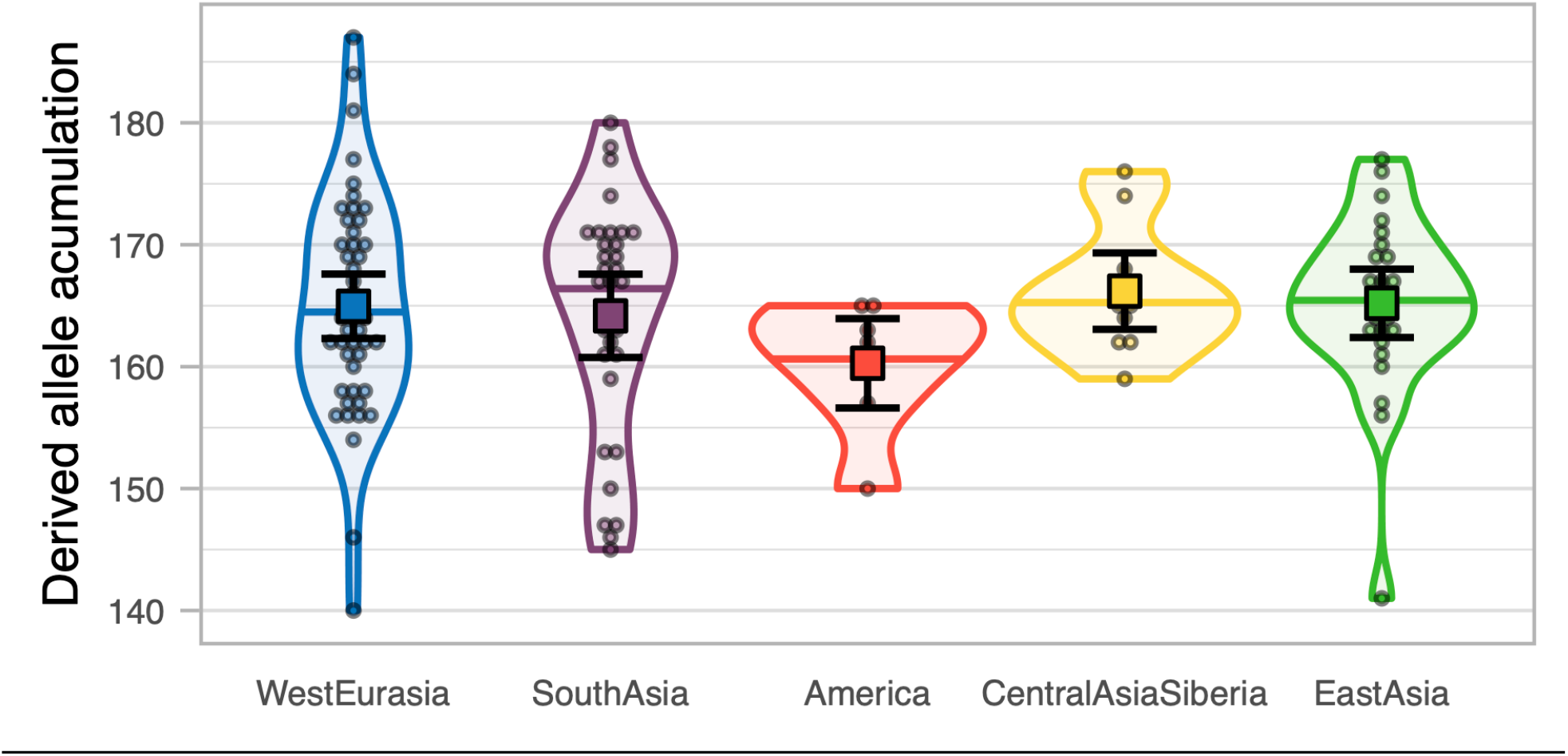
Mean derived allele accumulation of the 7-mutation types per region in the Y Chromosome. **a)** The mean number of derived alleles of each mutation type accumulated among individuals of the 5 regions (colour coded). The 95%CI of each mean is shown as error bars. **b)** The number of derived alleles of each mutation type per region (colour coded) as violin plot. Individual values are shown as dots. The median is shown as a horizontal line in each violin plot. The mean and its 95%CI of each distribution is shown as a coloured square with their corresponding error bars.

**Table S9.** **Derived allele accumulation per region for the X chromosome in female individuals**. Summary statistics of the derived allele accumulation per region on the X chromosome of females. For each region, the mean and the of SE (<u>S1</u>) is provided.

| Region | Number of samples | Derived allele accumulation (X chromosome) |  |
| --- | --- | --- | --- |
|  |  | mean | SE |
| West Eurasia | 23 | 2,820.21 | 21.17 |
| South Asia | 8 | 2,911.41 | 26.69 |
| America | 13 | 2,839.21 | 21.73 |
| Central Asia Siberia | 16 | 2,818.77 | 18.56 |
| East Asia | 20 | 2,900.35 | 17.46 |

**Table S10.** **Derived allele accumulation per region for the Y chromosome in male individuals**. Summary statistics of the derived allele accumulation per region on the X chromosome of males. For each region, the mean and the of SE (<u>S1</u>) is provided.

| Region | Number of samples | Derived allele accumulation (Y chromosome) |  |
| --- | --- | --- | --- |
|  |  | mean | SE |
| West Eurasia | 45 | 164.95 | 1.35 |
| South Asia | 31 | 164.17 | 1.74 |
| America | 7 | 160.28 | 1.87 |
| Central Asia Siberia | 10 | 166.20 | 1.60 |
| East Asia | 25 | 165.20 | 1.43 |

## 11 Datasets

Data1_archaicfragments.txt: Archaic fragments found in individuals from the 5 main geographical regions and ancient samples in the SGDP investigated in this study. Each line is a fragment with the following attributes:

1. name: individual the fragment belongs to.
2. region: region that the individual belongs to as defined by ^3^.
3. chrom: chromosome in which the fragment is located.
4. start: starting fragment position in hg19 coordinates.
5. end: ending fragment position in hg19 coordinates.
6. length: fragment length (end -start).
7. MeanProb: mean posterior probability for the fragment outputted by the ^11^ method.
8. snps: number of SNPs found in the fragment that are not segregating in any of the Sub Saharan African genomes (S3).
9. Altai: number of SNPs found in the fragment that are shared with the Altai Neanderthal ^33^.
10. Denisova: number of SNPs found in the fragment that are shared with the Denisova ^34^.
11. Vindija: number of SNPs found in the fragment that are shared with Vindija Neanderthal ^12^.

Data2_mutationspectrum.txt: Counts of derived alleles classified into the 96 mutation types for the extant samples of the SGDP, per chromosome. Each line has the following attributes:

1. ind: individual identifier
2. reg: region that the individual belongs to as defined by ^3^.
3. sex: individual sex defined by ^3^. M = male, F = female.
4. chrom: chromosome which the counts belong to.
5. fiv: contiguous 5*’* base pair of the focal SNP
6. anc: ancestral allele of the mutation
7. thr: contiguous 3*’* base pair of the focal SNP
8. der: ancestral allele of the mutation
9. counts: number of mutation types found

